# Phosphorylation by casein kinase 2 ensures ER-phagy receptor TEX264 binding to ATG8 proteins

**DOI:** 10.1101/2022.02.11.480038

**Authors:** Haruka Chino, Akinori Yamasaki, Koji L Ode, Hiroki R Ueda, Nobuo N Noda, Noboru Mizushima

## Abstract

Selective autophagy cargos are recruited to autophagosomes primarily by interacting with autophagosomal ATG8 family proteins via the LC3-interacting region (LIR). The upstream sequence of most LIRs contains negatively charged residues such as Asp, Glu, and phosphorylated Ser and Thr. However, the significance of LIR phosphorylation (compared with having acidic amino acids) and the structural basis of phosphorylated LIR–ATG8 binding are not entirely understood. Here, we show that the serine residues upstream of the core LIR of the endoplasmic reticulum (ER)-phagy receptor TEX264 are phosphorylated by casein kinase 2, which is critical for its interaction with ATG8s, autophagosomal localization, and ER-phagy. Structural analysis showed that phosphorylation of these serine residues increased binding affinity by producing multiple hydrogen bonds with ATG8s that cannot be mimicked by acidic residues. This binding mode is different from those of other ER-phagy receptors that utilize a downstream helix, which is absent from TEX264, to increase affinity. These results suggest that phosphorylation of the LIR is critically important for strong LIR–ATG8 interactions, even in the absence of auxiliary interactions.

## Introduction

Macroautophagy (hereafter, autophagy) is an intracellular degradation system through which cytoplasmic components are degraded in lysosomes. During autophagy, a portion of the cytoplasm is sequestered by autophagosomes, which then fuse with lysosomes to degrade their internal material (Nakatogawa, 2020; Søreng *et al*, 2018). Autophagy degrades intracellular materials not only randomly but also selectively (Gatica *et al*, 2018; Johansen & Lamark, 2020). In selective autophagy, certain soluble proteins, protein aggregates, organelles (e.g., mitochondria, endoplasmic reticulum (ER), and lysosomes), and intracellular pathogens are selectively recognized and degraded (Anding & Baehrecke, 2017; Chino & Mizushima, 2020; Gomes & Dikic, 2014; Palikaras *et al*, 2018). Selective autophagy is important for maintaining cellular homeostasis and is associated with human diseases (Dikic & Elazar, 2018; Hübner & Dikic, 2020; Mizushima & Levine, 2020). Most of the selective substrates are recognized by autophagosomal ATG8 family proteins directly or indirectly via selective autophagy receptors (Kirkin & Rogov, 2019). ATG8 proteins are ubiquitin-like proteins that can be covalently conjugated to phosphatidylethanolamine in the autophagic membrane (Mizushima, 2020). ATG8 proteins are classified into two subfamilies in mammals, namely, the LC3 (including LC3A, LC3B, and LC3C) and GABARAP (including GABARAP, GABARAPL1, and GABARAPL2) subfamilies.

Interactions with ATG8 proteins involve the LC3-interacting region (LIR) in selective substrates (Johansen & Lamark, 2020). The canonical LIR motif consists of the consensus sequence Θ_0_-X_1_-X_2_-Γ_3_ (called the core LIR), with Θ representing an aromatic residue (W/F/Y), Γ an aliphatic residue (L/V/I), and X any amino acid. The side chains of the aromatic and aliphatic residues of the LIR motif interact with two hydrophobic pockets on the surface of ATG8s to stabilize an intermolecular β-sheet formed by the LIR motif and β-strand 2 within ATG8s (Suzuki *et al*, 2014). In addition to these canonical interactions, other interactions sometimes critically enhance binding. For example, negatively charged residues are often found at the X_-3_, X_-2_, X_-1_, X_1_, and X_2_ positions. Acidic residues such as Asp and Glu and phosphorylated Ser and Thr at X_-3_, X_-2_, and X_-1_ can form electrostatic interactions with Lys46 and Lys48 of Atg8, whereas acidic residues at X_1_ and X_2_ form electrostatic interactions with Arg67 and Arg28 of Atg8 in yeasts (Noda *et al*, 2010). Phosphorylation of Ser and Thr residues at these positions could serve as a molecular switch. Upon bacterial infection, microbe-derived lipopolysaccharides induce phosphorylation of S177 (X_-1_) of optineurin by TANK binding kinase 1 (TBK1) and increase the affinity to ATG8s, promoting selective autophagy (Rogov *et al*, 2013; Wild *et al*, 2011). In addition, the mitophagy receptors BNIP3 and NIX (also known as BNIP3L) are also phosphorylated to function (Rogov *et al*, 2017; Zhu *et al*, 2013). These publications indicated the formation of additional intermolecular hydrogen bonds between Ser in positions X_-1_ and X_-2_ and Arg10, Arg11, Lys51 in LC3/GABARAPs. Besides these, some LIRs possess an α-helix downstream of the core LIR, which markedly increases their affinity with ATG8s through the additional intermolecular hydrogen bonds between electronegative residues after the core LIR and α-helix 3 of ATG8s (R69 and R70 in LC3B) (Johansen & Lamark, 2020; Li *et al*, 2018; Mochida *et al*, 2020; Sakurai *et al*, 2017). However, the structural basis by which these residues surrounding the core LIR enhance binding to ATG8s is not completely understood because these previous reports utilized nuclear magnetic resonance analysis or crystallography using phosphomimicking peptides or single-chain constructs.

In this study, we show that the serine residues upstream of the core LIR of TEX264, a receptor for autophagic degradation of the endoplasmic reticulum (ER-phagy) (An *et al*, 2019; Chino *et al*, 2019), are phosphorylated by casein kinase 2 (CK2). Phosphorylation of the LIR was required for the interaction of TEX264 with ATG8s, autophagosomal localization, and ER-phagy receptor function. According to the results of crystallographic analysis, we propose that the phosphorylation of residues upstream of the core LIR is critically important to produce strong LIR–ATG8 interactions, even in the absence of auxiliary interactions, by producing multiple hydrogen bonds in addition to ionic interactions.

## Results

### Ser266, Ser269, Ser271, and Ser272 upstream of the core LIR in TEX264 are phosphorylated

TEX264 contains multiple highly conserved serine residues upstream of the core LIR (Fig 1A). An anti-phosphoserine antibody reacted to immunopurified FLAG-tagged TEX264 (TEX264–FLAG), suggesting that the serine residues of TEX264 were phosphorylated (Fig 1B). To determine the exact phosphorylation sites, we performed tandem mass spectrometry (MS/MS). To detect a peptide containing the LIR sequence in MS/MS analysis, we generated a TEX264^G280K^ mutant such that the otherwise too long trypsinized peptides around the LIR in wild-type (WT) TEX264 (SEHSYSESGASGSSFEELDLEGEGPLGESR) could be digested by trypsin at the introduced Lys residue. MS/MS analysis revealed multiple phosphorylations in various combinations around the LIR sequence. Among them, Ser266, Ser269, Ser271, and Ser272 were identified as being phosphorylated (Fig 1A, asterisks, Fig EV1A)

**Figure 1.**
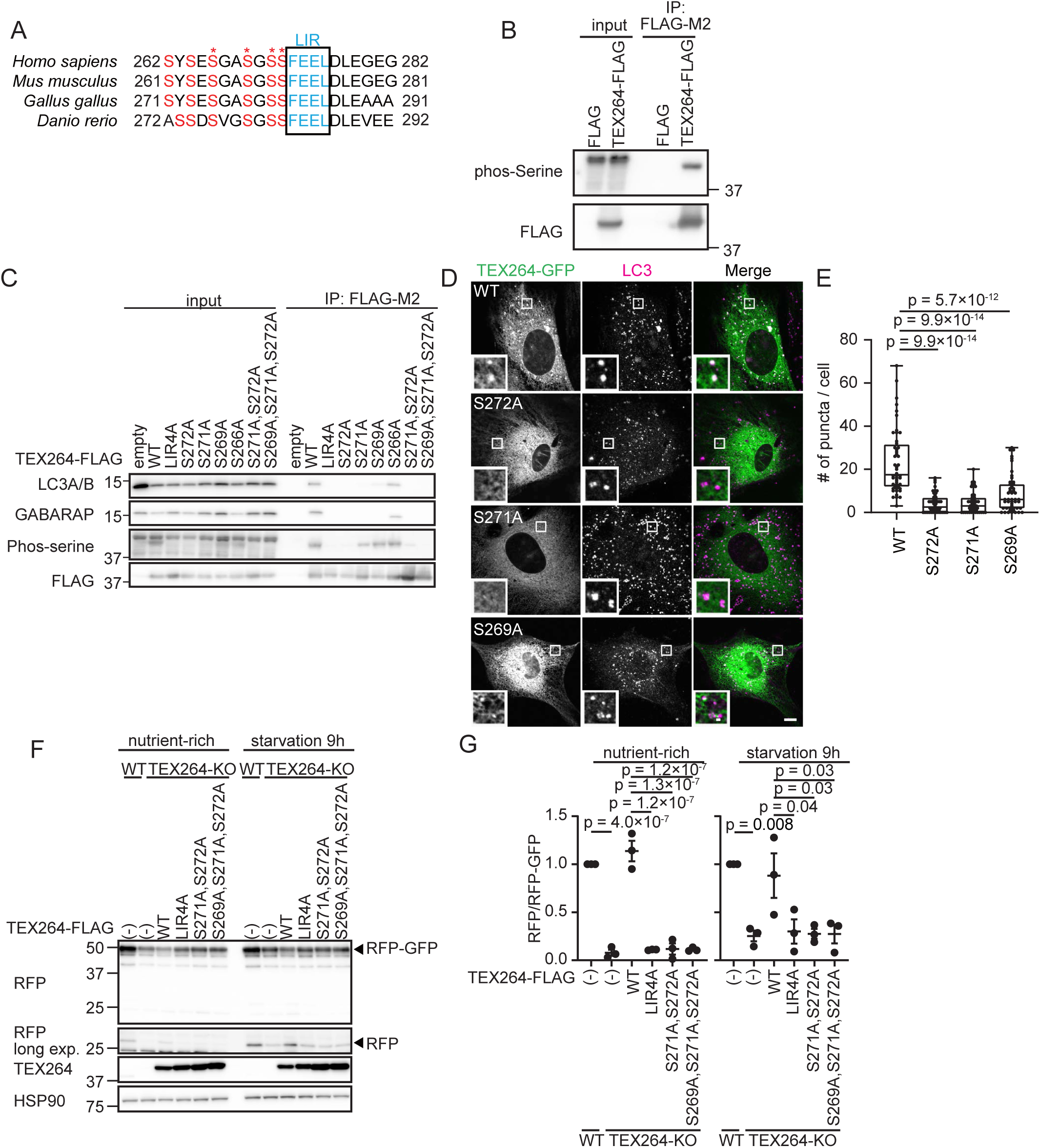
Phosphorylation of Ser271 and Ser272 upstream of the TEX264 LIR is required for ATG8 binding and autophagosomal localization. (A) Alignment of the LIR motif of TEX264 homologs in vertebrates. Asterisks show the phosphorylation sites identified by mass spectrometry. (B) HEK293T cells transiently expressing FLAG and FLAG-tagged TEX264 were subjected to immunoprecipitation (IP). Inputs (20% of the lysates) and immunoprecipitates (from 80% of the lysates) were analyzed by immunoblotting using the indicated antibodies. (C) HEK293T cells transiently expressing WT or mutated TEX264-FLAG were subjected to immunoprecipitation with an anti-FLAG antibody and detected with the indicated antibodies. (D and E) MEFs stably expressing TEX264-GFP or its mutant were cultured in starvation media and immunostained with an anti-LC3 antibody. Bars: 10 µm and 1 µm (insets) (D). Quantification of the number of TEX264 puncta per cell. Solid bars indicate the medians, boxes the interquartile range (25th to 75th percentile), and whiskers the 0th to 100th percentile. Differences were statistically analyzed by one-way ANOVA and Tukey’s multiple comparison test. Data were collected from 48 cells for each cell type (E). (F and G) WT and TEX264-KO HeLa cells (with or without the indicated TEX264 mutants with a C-terminal FLAG tag) stably expressing the ER-phagy reporter (ss-RFP-GFP-KDEL) were cultured in the presence of doxycycline for 24 h to induce the reporter. After doxycycline was removed, cells were cultured in starvation medium lacking amino acids and serum for 9 h (F). The band intensities of RFP and RFP-GFP were quantified and the ratio of RFP:RFP-GFP (normalized to WT) is shown. Data represent the mean ± standard error of the mean (SEM) of three independent experiments. Differences were statistically analyzed by one-way ANOVA and Tukey’s multiple comparison test (G).

### Ser271 and Ser272 are essential for ATG8 binding and autophagosomal localization of TEX264

To investigate the importance of the phosphorylation of Ser266, Ser269, Ser271, and Ser272 in TEX264, we replaced them with alanine and tested the interaction of TEX264 with ATG8s. Among mammalian ATG8s, TEX264 is known to interact with LC3A, LC3B, GABARAP, and GABARAPL1 (An *et al*., 2019; Chino *et al*., 2019). Alanine-substitution mutants of either or any combination of the serine residues decreased the level of serine phosphorylation (Fig 1C). Unexpectedly, mutation of the core LIR (LIR4A, FEEL/AAAA) also impaired serine phosphorylation (discussed later). As with the core LIR mutant, single or multiple serine residue mutants including Ser271 and Ser272 abolished the interaction with LC3A/B and GABARAP, although TEX264^S269A^ still weakly bound to GABARAP (Fig 1C). Alanine substitution of Ser266 did not affect the interaction with LC3A/B and GABARAP (Fig 1C). C-terminal GFP-tagged TEX264 (TEX264-GFP) was present in the ER and formed punctate structures with LC3 during starvation (Fig 1D and E). By contrast, although the TEX264^S269A^, TEX264^S271A^, and TEX264^S272A^ mutants still localized to the ER, their accumulation at LC3 structures was significantly suppressed (Fig 1D and E). Consistent with the immunoprecipitation data, TEX264^S266A^ formed punctate structures and localized to autophagosomes like wild-type TEX264 (Fig EV1B and C). These results suggest that the serine residues, particularly Ser271 and Ser272, upstream of the core LIR are required for ATG8 binding and autophagosomal localization of TEX264.

To determine whether these serine residues are important for the ER-phagy receptor function of TEX264, we monitored ER-phagy activity using a doxycycline-inducible ER-phagy reporter with an N-terminal ER signal sequence followed by tandem monomeric RFP and GFP and the ER retention sequence KDEL (Chino *et al*., 2019). Upon delivery of this reporter to lysosomes, GFP, but not RFP, is degraded, producing an RFP fragment. When expressed in TEX264-knockout (KO) cells, WT TEX264-FLAG recovered ER-phagy activity, but the TEX264^LIR4A^, TEX264^S271A, S272A^, and TEX264^S269A, S271A, S272A^ mutants did not (Fig 1F and G). Thus, Ser269, Ser271 and Ser272 are required for the ER-phagy receptor function of TEX264.

### Phosphorylated TEX264 localizes to autophagosomes

To further characterize the phosphorylation of TEX264, we generated an antibody against the peptide GASGpSpSFEEL (a.a. 267–276 of TEX264) containing phosphorylated Ser271 and Ser272 (pTEX264), because phosphorylation of Ser271 and Ser272 is particularly important for its interaction with ATG8s (Fig1C). The anti-pTEX264 antibody could detect endogenous and exogenous TEX264, but not the exogenous TEX264^S271,272A^ mutant (Fig EV1D). Phosphatase treatment completely abolished the pTEX264 signals (Fig EV1E), validating that the anti-pTEX264 antibody specifically recognizes phosphorylated TEX264.

Although this antibody barely detected endogenous TEX264 by immunofluorescence microscopy, it could detect stably overexpressed TEX264-GFP (Fig 2A). The GFP signal of TEX264–GFP was observed as reticular and punctate patterns, whereas the pTEX264 signal mainly accumulated at autophagosomes under starvation conditions. The number of pTEX264-positive puncta increased following the treatment with the lysosome inhibitor bafilomycin A_1_, indicating that pTEX264 is efficiently degraded in lysosomes (Fig 2A and B). We determined whether endogenous pTEX264 localized to autophagosomes by a membrane floatation assay. In a stepwise OptiPrep gradient, autophagosomes and autolysosomes containing LC3-II and LAMP1 were collected in fractions 3 and 5, where endogenous pTEX264 was cofractionated (Fig 2C). The ER marker SEC61B was not enriched in these fractions. These data suggest that phosphorylated TEX264 preferentially localizes to autophagosomes under autophagy-induction conditions such as starvation and mTORC1 inhibition.

**Figure 2.**
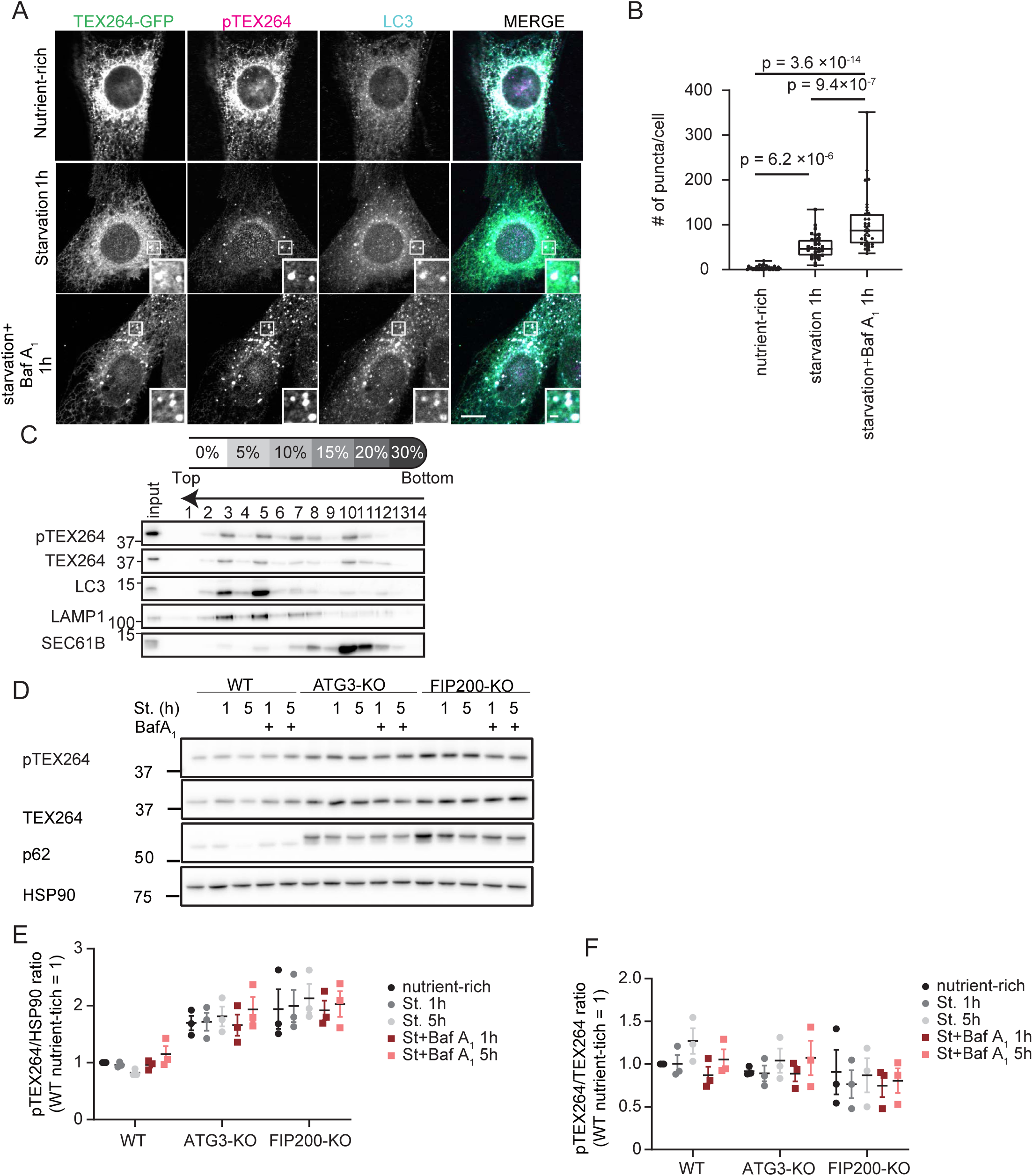
Phosphorylated TEX264 localizes to autophagosomes. (A and B) MEFs stably expressing TEX264-GFP were cultured in starvation media with or without bafilomycin A_1_ (Baf A_1_) and immunostained with anti-pTEX264 and anti-LC3 antibodies. Bars: 10 µm and 1 µm (insets) (A). Quantification of the number of pTEX264 puncta per cell. Solid bars indicate the medians, boxes the interquartile range (25th to 75th percentile), and whiskers the 0th to 100th percentile. Differences were statistically analyzed by one-way ANOVA and Tukey’s multiple comparison test. Data were collected from 35 cells for each cell type (B). (C) HeLa cells were treated with 250 nM Torin 1 and Baf A_1_ for 16 h, and cell homogenates were subjected to an OptiPrep membrane flotation analysis. (D, E and F) WT, ATG3-KO, and FIP200-KO HeLa cells were cultured in starvation media lacking amino acids and serum with or without Baf A_1_ for 1 or 5 h. Cell lysates were analyzed by immunoblotting using the indicated antibodies (D). Relative changes of the ratio of band intensities of phosphorylated TEX264 to HSP90 and total TEX264 during starvation are shown (E and F). Data represent the mean ± SEM of three independent experiments.

Consistently, the level of phosphorylated TEX264 was decreased under starvations, which was restored by lysosome inhibition, and increased in autophagy-deficient ATG3-KO and FIP200-KO cells (Fig 2D and E). However, the ratio of phosphorylated TEX264 to total TEX264 was not changed significantly under these conditions (Fig 2D and F). These data suggest that, while phosphorylated TEX264 is efficiently recruited to autophagosomes, there is an equilibrium between the phosphorylated and dephosphorylated forms.

### CK2 is important for starvation-induced ER-phagy through TEX264 phosphorylation

Ser271 and Ser272 of TEX264 fulfill the criteria for a consensus motif recognized by CK2, namely, S-X-X-E/D/Yp/Sp (Meggio *et al*, 1994; Pinna, 2002; Rekha & Srinivasan, 2003). Thus, we tested the requirement of CK2 activity for TEX264 phosphorylation by using CX4945, a specific inhibitor of CK2. CX4945 suppressed TEX264 phosphorylation, whereas a CK1 inhibitor (D4476) did not (Fig 3A). Also in an invitro system, recombinant TEX264-FLAG was phosphorylated by CK2, which was inhibited by the CK2 inhibitor (Fig 3B). These results suggest that CK2 directly phosphorylates TEX264. The association between TEX264 and all the three isoforms of CK2A (catalytic subunits of CK2) was shown using an immunoprecipitation assay (Fig 3C).

**Figure 3.**
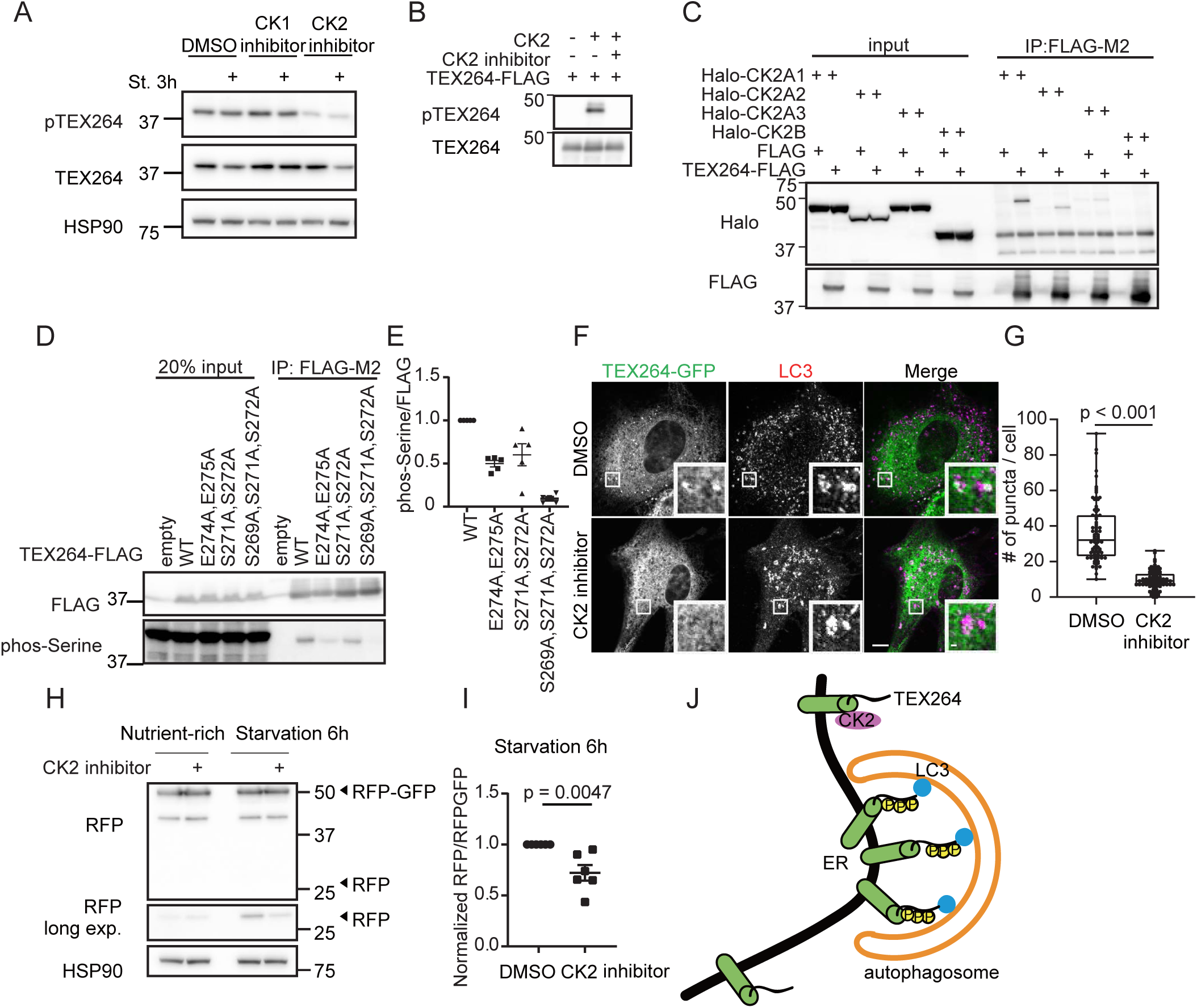
CK2 is important for starvation-induced ER-phagy through TEX264 phosphorylation. (A) HeLa cells were cultured with 20 nM CK1 inhibitor (D4476) or CK2 inhibitor (CX4945) for 3 h, followed by starvation (with the inhibitors) for 3 h. Cell lysates were analyzed by immunoblotting using the indicated antibodies. (B) Recombinant TEX264-FLAG was incubated with or without CK2 and the CK2 inhibitor (CX4945) in the presence of 200 µM ATP for 1 h. (C) HEK293T cells transiently expressing TEX264-FLAG and one of the three Halo-CK2A isoforms or Halo-CK2B were subjected to immunoprecipitation with an anti-FLAG antibody and detected with anti-Halo and anti-FLAG antibodies. (D and E) HEK293T cells transiently expressing WT or TEX264-FLAG mutants were subjected to immunoprecipitation (IP). Inputs (20% of the lysates) and immunoprecipitates (from 80% of the lysates) were analyzed by immunoblotting using the indicated antibodies (D). The band intensities in the IP fractions were quantified, and the phospho-serine:FLAG ratio was calculated. Data represent the mean ± SEM of five independent experiments. Differences were statistically analyzed by one-way ANOVA and Tukey’s multiple comparison test (E). (F and G) MEFs stably expressing TEX264-GFP were cultured with the CK2 inhibitor for 3 h followed by starvation (with the CK2 inhibitor) and immunostained with an anti-LC3 antibody. Scale bars represent 10 μm and 1 μm (insets) (F). Quantification of the number of TEX264 puncta per cell. Solid bars indicate the medians, boxes the interquartile range (25th to 75th percentile), and whiskers the 0th to 100th percentile. Differences were statistically analyzed by an unpaired two-tailed Student’s t-test. Data were collected from 85 cells for each cell type (G). (H and I) WT HeLa cells stably expressing the ER-phagy reporter were cultured in the presence of doxycycline for 24 h to induce the reporter. After doxycycline was removed, cells were cultured with 20 nM CK2 inhibitor for 3 h followed by starvation (with the CK2 inhibitor) for 6 h (H). The band intensities of RFP and RFP-GFP were quantified and the ratio of RFP:RFP-GFP (normalized to WT) is shown. Data represent the mean ± SEM of six independent experiments. Differences were statistically analyzed by an unpaired two-tailed Student’s t-test (I). (J) Model of the CK2-mediated phosphorylation of TEX264 in ER-phagy.

Since the two glutamate residues (Glu274, Glu275) inside the core LIR are part of the CK2 consensus motif, they may be required for the first priming event, which triggers processive phosphorylation of further upstream serine residues (Zhou *et al*, 2017). To test this hypothesis, we analyzed the phosphorylation status of an alanine substitution mutant of TEX264 (TEX264^E274A,E275A^). We used the anti-phosphoserine antibody instead of the anti-pTEX264 antibody because it could not detect the TEX264^E274A,E275A^ mutant regardless of its phosphorylation status. The TEX264^E274A,E275A^ mutant as well as the TEX264^S271A, S272A^ and TEX264^S269A,S271A,S272A^ mutants showed significantly reduced levels of serine phosphorylation (Fig 3D and E). This also explained why the core LIR4A mutant significantly decreased the level of serine phosphorylation (Fig 1C). These results suggest that the two glutamates inside the core LIR are the priming site for CK2-mediated TEX264 phosphorylation.

To determine the significance of the CK2-mediated phosphorylation of TEX264, we analyzed the effect of the CK2 inhibitor CX4945 on the autophagosomal localization of TEX264. CK2 inhibition significantly impaired the formation of TEX264 puncta (Fig 3F and G), suggesting that CK2-mediated phosphorylation is important for the autophagosomal localization of TEX264. Next, we measured ER-phagy activity using the ER-phagy reporter and found that the CK2 inhibitor significantly suppressed the production of the RFP fragment under starvation conditions (Fig 3H and I). The CK2 inhibitor did not inhibit starvation-induced bulk autophagy judged using the GFP-LC3-RFP reporter (Fig EV2) (Kaizuka *et al*, 2016). These data suggest that CK2 activity is important for starvation-induced ER-phagy through TEX264 phosphorylation (Fig 3J).

### Phosphorylation of the TEX264 LIR increases its affinity to ATG8s

Next, we determined whether the phosphorylation of the LIR in TEX264 promoted binding to ATG8s by an isothermal titration calorimetry (ITC) assay (Fig 4A, Fig EV3A and Table EV1). The TEX264 peptide (residues 269–278) with S272 phosphorylation (p-S272) showed enhanced affinity to LC3B and GABARAP (37- and 23-fold, respectively). Moreover, multiple phosphorylations (p-S271, p-S272 and p-S269, p-S271, p-S272) further strengthened the affinity of TEX264 to LC3B and GABARAP, resulting in sub-µM Kd values (0.25–0.52 µM). S266 phosphorylation (p-S266) only slightly enhanced the affinity to LC3B and GABARAP (1.8- and 6.5-fold, respectively) (Fig EV3B). The phosphomimetic TEX264^S271D, S272D^ (2SD-LIR), TEX264^S269D, S271D, S272D^ (3SD-LIR), TEX264^S271, 272E^ (2SE-LIR), and TEX264^S269E, S271E, S272E^ (3SE-LIR) peptides also demonstrated increased binding to LC3B and GABARAP compared with WT TEX264 and TEX264^S271A, S272A^ peptides; however, the Kd value was within the order of µM (the minimum was 5.0 and 2.1 µM for LC3B and GABARAP, respectively). Consistently, compared with WT TEX264, _TEX264_S269D, S271D, S272D _and TEX264_S269E, S271E, S272E _showed a significant_ reduction in their affinity to LC3A, LC3A/B (the antibody reacts to both LC3A and LC3B), and GABARAP (Fig 4B), accumulation at autophagosomes during starvation (Fig 4C and D), and ability to promote ER-phagy (Fig 4E and F). These data suggest that multiple phosphorylations of the upstream region of the core LIR sufficiently strengthen the affinity of TEX264 to ATG8s to mediate ER-phagy, which cannot be complemented by Asp or Glu mutations.

**Figure 4.**
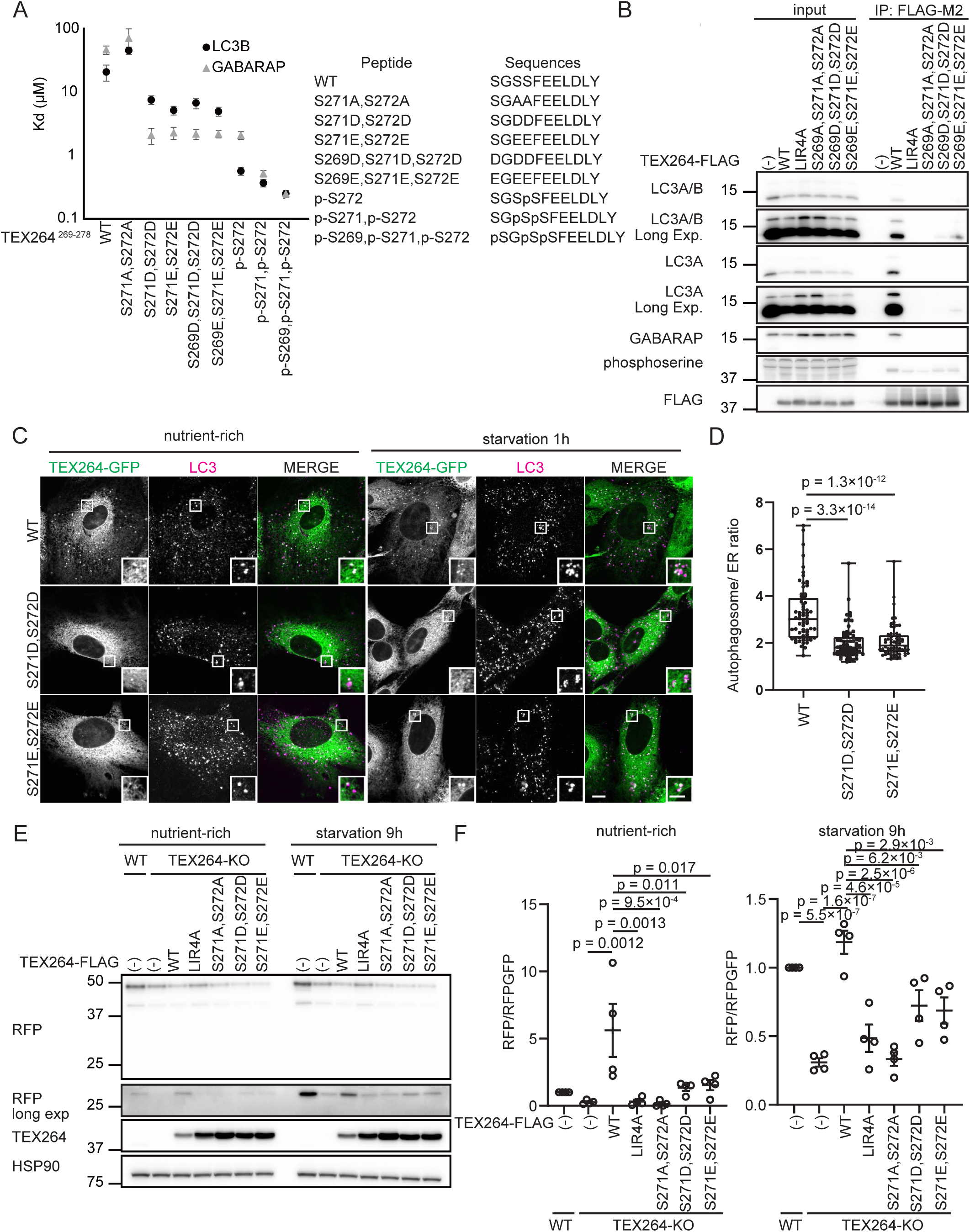
Phosphorylation of TEX264 LIR increases its affinity to ATG8s. (A) Binding affinity of TEX264 LIR peptides (TEX264^269–278^) to LC3 and GABARAP was measured by ITC. The error bars show the fitting error. (B) HEK293T cells transiently expressing WT or mutated TEX264-FLAG were subjected to immunoprecipitation with an anti-FLAG antibody and detected with the indicated antibodies. (C and D) MEFs stably expressing TEX264-GFP or its mutants were cultured in starvation media and immunostained with an anti-LC3 antibody. Bars: 10 µm and 5 µm (insets) (C). The intensity of the GFP signal under starvation conditions was quantified at TEX264-GFP puncta and the ER. The puncta:ER signal ratio was calculated. Solid bars indicate the medians, boxes the interquartile range (25th to 75th percentile), and whiskers the 0th to 100th percentile. Differences were statistically analyzed by one-way ANOVA and Tukey’s multiple comparison test. Data were collected from 70 puncta for each cell type (D). (E and F) WT and TEX264-KO (expressing WT or TEX264-FLAG mutants) HeLa cells stably expressing the ER-phagy reporter were cultured in the presence of doxycycline for 24 h to induce the reporter. After doxycycline was removed, the cells were cultured in starvation medium lacking amino acids and serum for 9 h (E). The band intensities of RFP and RFP-GFP were quantified and the ratio of RFP:RFP-GFP (normalized to WT) is shown. Data represent the mean ± SEM of four independent experiments. Differences were statistically analyzed by one-way ANOVA and Tukey’s multiple comparison test (F).

### Crystallographic analysis reveals canonical interactions between GABARAP and TEX264 LIR

To elucidate the structural basis of ATG8 binding by TEX264 LIR and its enhancement by phosphorylation, we determined the crystal structure of GABARAP complexed with phosphorylated TEX264 LIR (residues 269–278 with p-S271,p-S272; referred to as LIR^p-S271,p-S272^) and that of GABARAP fused to TEX264 LIR with the S272D mutation (residues 271–281, referred to as LIR^S272D^) at a resolution of 3.0 and 2.0 Å, respectively (Fig 5A and B, Table EV2, Protein Data Bank [PDB] code 7VEC and 7VED, respectively). The asymmetric unit contained 12 and 2 copies of GABARAP for LIR^p-S271,p-S272^ and LIR^S272D^, respectively, which formed a complex with each LIR in a similar manner (Fig EV4A–D). In both structures, the TEX264 LIR formed an intermolecular β-sheet with β-strand 2 and bound to the hydrophobic pockets 1 and 2 of GABARAP using the side-chains of Phe273 at position Θ_0_ and Leu276 at position Γ3, respectively, as with canonical LIRs (Fig 5A and B). Moreover, Glu274 at position X_1_ and Glu275 at position X_2_ and/or Asp277 at position X4 formed electrostatic interactions with Arg67 and Arg28 of GABARAP, respectively. In the case of LIR^p-S271,p-S272^, Leu278 at position X_5_ formed hydrophobic interactions with α-helix 3 of GABARAP (Fig 5A). These auxiliary interactions within the core LIR and its downstream region are also observed for other LIRs to enhance their binding affinity to ATG8s. These observations suggest that the manner by which the TEX264 LIR interacts with ATG8s is similar to that of other canonical LIRs.

**Figure 5.**
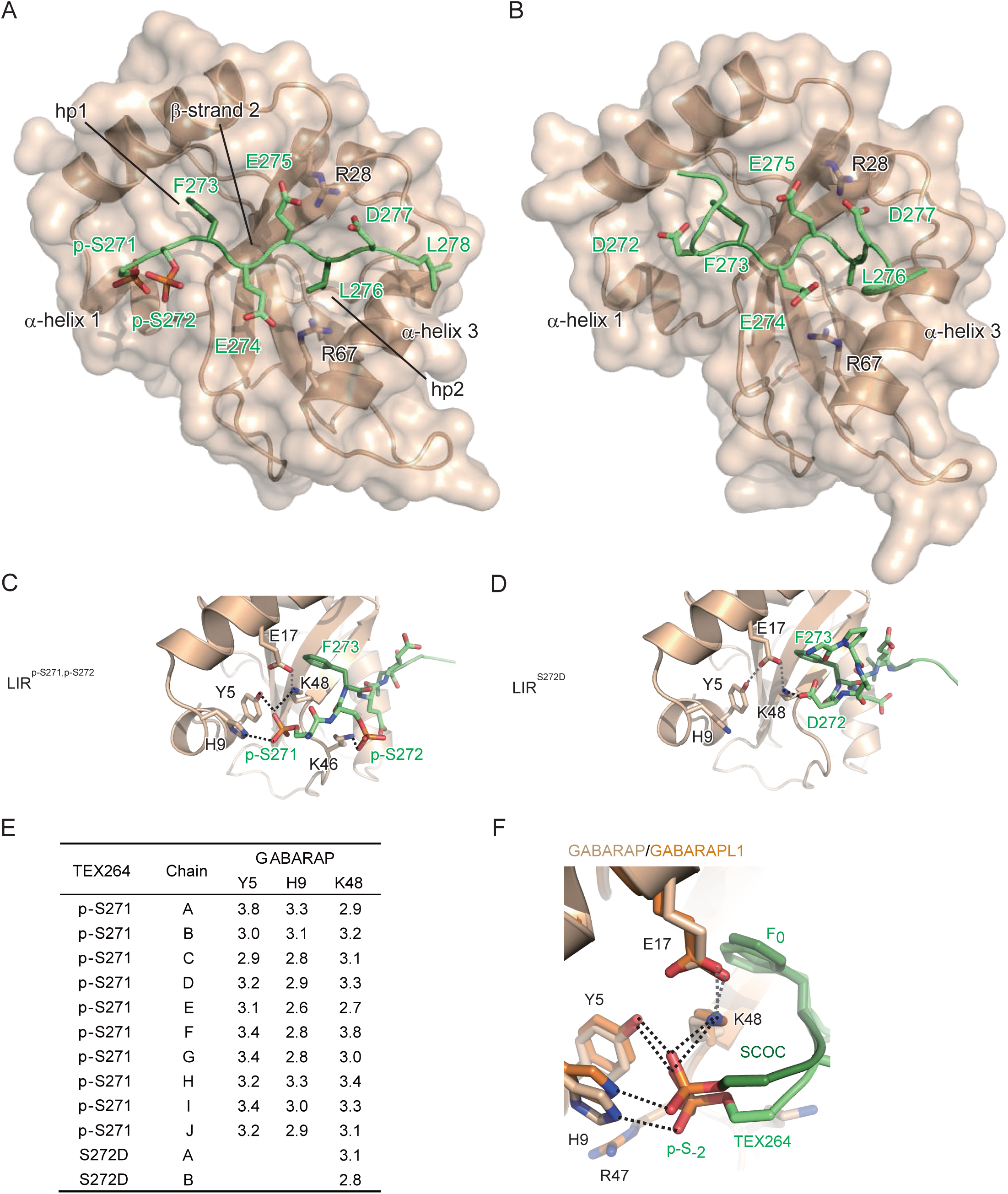
Phosphorylation of the LIR in TEX264 produces hydrogen bonds to interact with GABARAP. (A) Crystal structure of GABARAP complexed with TEX264^269–278^ peptide containing p-Ser271 and p-Ser272. hp1 and hp2 indicate hydrophobic pockets 1 and 2, respectively. TEX264 peptide is colored green. (B) Crystal structure of GABARAP fused with TEX264^271–281^ peptide with the S272D mutation. TEX264 peptide is colored green. (C and D) Close-up view of the interaction between GABARAP and the upstream region of LIR^p-S271,p-S272^ (C) or LIR^S272D^ (D). Broken lines indicate possible hydrogen bonds between GABARAP and LIR (colored back) and within GABARAP (colored gray). (E) Summary of the distances (Å) of possible hydrogen bonds mediated by Tyr5, His9, and Lys48 of GABARAP with TEX264 LIR. (F) Structural comparison between the GABARAP-TEX264 LIR^p-S271,p-S272^ complex and GABARAPL1-SCOC LIR complex (PDB 7AA7). Possible hydrogen bonds are shown as in C and D. F_0_ and p-S-_2_ indicate Phe located at position Θ_0_ and phosphorylated Ser located at position X_-2_, respectively.

### Phosphorylation at the X_-2_ position in TEX264 LIR produces multiple hydrogen bonds for strong interactions with GABARAP

Although the conformation and interaction manner of the core LIR motif with GABARAP are essentially similar between the LIR^p-S271,p-S272^ and LIR^S272D^ structures, those of the upstream region of the core LIR are totally distinct between them (Fig 5C and D). In the LIR^S272D^ structure, the main-chain of LIR was bent at Asp272 (position X_-1_) at a right angle and the upstream region was located away from α-helix 1 of GABARAP. The side-chain of Asp272 formed a hydrogen bond with that of GABARAP Lys48. On the other hand, in the LIR^p-S271,p-S272^ structure, the upstream region had a stretched β-conformation continuous with the core LIR motif and interacted with α-helix 1 of GABARAP. The phosphate group of p-Ser272 (position X_-1_) formed a hydrogen bond with the side-chain of GABARAP Lys46, while that of p-Ser271 (position X_-2_) formed three hydrogen bonds with the side-chain of Tyr5 and His9 at α-helix 1 and Lys48 (whose side-chain is locked by Glu17) of GABARAP (Fig 5E). We confirmed the contribution of Tyr5, His9, and Lys48 to the high affinity with TEX264 LIR p-Ser271,p-Ser272 by ITC. The Kd values of GABARAP K48A and Y5F H9A mutants were ∼2 µM (Fig EV3A), which were comparable with that between GABARAP and TEX264 LIR with single phosphorylation at Ser272 (Fig 4A). Consistently, alanine substitution of Tyr5, His9, and Lys48 in GABARAP significantly reduced the affinity to TEX264 in vivo (Fig EV4E and F). These results suggest that the 4-fold enhancement of the affinity of TEX264 with GABARAP by phosphorylation at Ser271 (p-Ser272 vs. p-Ser271,p-Ser272 in Fig 4A), which enables sub-µM affinity, is presumably established by these multiple hydrogen bonds.

We then wondered whether the formation of multiple hydrogen bonds of p-Ser at position X_-2_ is specific to TEX264 or conserved among other LIRs. In contrast to the strict conservation of Lys48 among all six ATG8 paralogs, His9 was conserved only among the GABARAP family and Tyr5 was conserved in GABARAP/GABARAPL1. We inspected all of the structures of GABARAP/GABARAPL1 deposited in the PDB and found that the crystal structure of GABARAPL1 complexed with the phosphorylated LIR from the Golgi protein short coiled-coil protein (SCOC) (PDB 7AA7; (Wirth *et al*, 2021)) showed a strikingly similar interaction: the phosphate group of p-Ser at position X_-2_ of SCOC LIR formed three hydrogen bonds with the side-chain of His9, Lys48, and either Tyr5 or Arg47 of GABARAPL1 (Fig 5F) (Wirth *et al*., 2021). The authors revealed that Ser phosphorylation at position X_-2_ of the long SCOC LIR peptide (containing four consecutive Glu residues at positions X-7–X-4) marginally enhanced binding affinity (2.4- and 2.6-fold for GABARAP and GABARAPL1, respectively), whereas phosphorylation at position X_-1_ did not. Moreover, they also showed that Ser phosphorylation at position X_-2_ of the short SCOC LIR peptide (lacking four Glu residues at positions X_-7_–X_-4_) enhanced binding affinity dramatically (67- and 275-fold with GABARAP and GABARAPL1, respectively) (Wirth *et al*., 2021). Our ITC data showed that two or three Asp/Glu mutations at positions X_-3_–X_-1_ or single phosphorylation at position X_-1_ of TEX264 LIR enhanced its binding affinity with GABARAP to a similar extent (approximately 20-fold), which was further enhanced (4-fold) by phosphorylation at position X_-2_ (Fig 4A). These observations suggest that the enhancement of binding affinity by phosphorylation at the upstream region of the core LIR is achieved by two distinct mechanisms: the first endows negative charges to the LIR for non-specific interactions with positively-charged ATG8s (Noda *et al*., 2010), and the second creates specific interactions with ATG8s, as observed for triple hydrogen bonding between p-Ser at X_-2_ and Tyr5, His9, and Lys48 of GABARAP/GABARAPL1. A phosphomimetic Asp or Glu mutation at the upstream region would be able to account for only the former mechanism since the chemical structure of the carboxyl side-chain of Asp/Glu cannot form similar specific hydrogen bonds with those observed for the phosphate group. Consistent with this idea, there is no example of triple hydrogen bonding of Asp or Glu to Tyr5, His9, and Lys48 of GABARAP/GABARAPL1 among the dozens of LIR-bound structures deposited in the PDB (Fig EV5). Thus, phosphorylation but not Asp/Glu incorporation at the upstream region of the core LIR is essential for TEX264 to generate ATG8-binding affinity strong enough (sub-µM K_d_ values) to initiate ER-phagy.

## Discussion

In this study, we showed that CK2-mediated phosphorylation of TEX264 is required for its ER-phagy receptor function, and the phosphorylated serine residues upstream of the core LIR are important for generating high affinity to ATG8s that cannot be mimicked by the substitution of acidic residues.

In many studies, phosphorylated Ser and Thr residues are substituted with acidic residues to mimic their phosphorylation status (Thorsness & Koshland, 1987). However, this approach does not always work. For example, inhibition of yeast Ste5, a MAPK cascade scaffold protein, requires multiple phosphorylations, but the replacement of Ser/Thr with Glu cannot mimic inhibition (Strickfaden *et al*, 2007). This may be because phosphorylated Ser and Thr are, nominally, charged twice as much as Glu and Asp at physiological pH (Cooper *et al*, 1983) and could, therefore, strengthen electrostatic interactions. However, it cannot fully explain the differences in binding. Our structural analysis suggests that the phosphorylated residue at position X_-2_ in TEX264 LIR could form three hydrogen bonds in an ideal configuration with GABARAP. This hydrogen-bonding pattern is quite similar to the recently reported binding mediated by p-Ser at position X_-2_ of SCOC LIR bound to GABARAPL1 (Wirth *et al*., 2021). In contrast, the interaction of the phosphorylated residue at position X_-1_ is weaker and less specific: p-Ser/p-Thr at position X_-1_ of TEX264 and SCOC LIRs formed only one and no hydrogen bond with GABARAP and GABARAPL1, respectively. These observations suggest that GABARAP and GABARAPL1 conserve the specific binding site (composed of Tyr5, His9, and Lys48) for the phosphorylated residue at position X_-2_ of LIRs, which plays a critical role in increasing binding affinity. Whether such a specific binding pocket is conserved among other members of ATG8s needs to be addressed in the future. LC3 family proteins have conserved Arg and Lys residues (Arg11 and Lys51 in the case of LC3B) that correspond to His9 and Lys48 of GABARAP, respectively, which might form hydrogen bonds and/or salt-bridges with p-S271 of TEX264 to enhance binding affinity. Collectively, we propose that phosphorylation of the LIR is important for not only spatiotemporal regulation but also strong interactions with ATG8s, which are accomplished by the formation of specific hydrogen bonds that cannot be mimicked by acidic residues, in addition to the previously described ionic interactions (Noda *et al*., 2010).

Some selective autophagy receptors have only acidic amino acids upstream of the core LIR. p62 (also known as SQSTM1), one of the most well-studied autophagy receptors, has a “DDD” sequence followed by the core LIR sequence “WTHL.” The Kd value of p62–LC3 interactions is 1.5 μM (Rozenknop *et al*, 2011), which is comparable with those of the interactions between TEX264 2D/2E-LIR and LC3B and between 3D/3E-LIR and LC3B. In contrast, other ER-phagy receptors such as FAM134B and SEC62 show high affinity to GABARAPs with Kd values of 0.11–0.30 μM and 0.19–0.57 μM, and to LC3s with Kd values of 0.21–1.11 μM and 0.92–5.18 μM, respectively (Mochida *et al*., 2020). These two receptors possess acidic LIRs with a C-terminal short helix, which is used in various LIRs to enhance ATG8-binding affinity to sub-μM or even smaller Kd values through interactions with α-helix 3. Although TEX264 LIR does not possess a C-terminal helix, multiple phosphorylated serine residues can instead generate high affinity (sub-μM Kd values) to ATG8s via specific interactions, which enable this receptor to initiate ER-phagy, which appears to require higher affinity with ATG8s than other types of autophagy.

Our data show that the ratio of phosphorylated TEX264 to total TEX264 is not changed under ER-phagy-inducing conditions. This may be because phosphorylated TEX264 is efficiently degraded and/or there is an equilibrium between the phosphorylated and dephosphorylated forms. To maintain the ratio, it is assumed that TEX264 phosphorylation is upregulated by starvation. It is also possible that phosphorylation could act as a switch as TEX264 has more than one function. Besides ER-phagy, TEX264 is required for DNA quality control at the inner nuclear membrane (Fielden *et al*, 2020). It is not known whether this nuclear function of TEX264 also requires phosphorylation. If not, phosphorylation and dephosphorylation could switch between these two different functions. This may be similar to the case of AMPK-mediated phosphorylation of the Beclin 1 LIR in response to nutrient starvation, which underlies the differential regulation of the pro- and non-autophagic functions of the VPS34 complex (Kim *et al*, 2013).

## Materials and Methods

### Cell culture

HeLa cells, HEK293T cells, and mouse embryonic fibroblasts (MEFs) were authenticated by RIKEN. They were cultured in Dulbecco’s modified Eagle’s medium (DMEM) (D6546; Sigma-Aldrich) supplemented with 10% fetal bovine serum (S1820-500; Biowest) and 2 mM L-glutamine (25030-081; Gibco) in a 5% CO2 incubator. For starvation treatment, cells were washed twice with phosphate-buffered saline (PBS) and incubated in amino acid-free DMEM (048-33575; Wako Pure Chemical Industries) without serum. FIP200-KO HeLa cells (Tsuboyama *et al*, 2016) and ATG3-KO HeLa cells (Sou *et al*, 2008) have been described previously. For bafilomycin A_1_ treatment, cells were cultured with 100 nM bafilomycin A_1_ (B1793; Sigma-Aldrich). For CK2 inhibitor treatment, cells were cultured with 20 µM CX4945 (S2248; Selleck). WT HeLa cells and TEX264-KO HeLa cells stably expressing the ER-phagy reporter pCW57.1-CMV-ssRFPGFP-KDEL (Chino *et al*., 2019) were cultured with 0.5 μg/mL doxycycline (D3447; Sigma-Aldrich) for 24 h. Thereafter, cells were washed with PBS twice and incubated in DMEM or amino acid-free DMEM without serum for 9 h.

### Plasmids

cDNA encoding human *TEX264* (Chino *et al*., 2019) was inserted into pMRX-IN (Morita *et al*, 2018) or pMRX-IB (Morita *et al*., 2018) (these plasmids were generated from pMXs (Kitamura *et al*, 2003)). DNAs encoding enhanced GFP and 3×FLAG were also used for tagging. Amino acid substitution constructs were prepared by PCR-mediated site-directed mutagenesis.

### Antibodies and reagents

For immunoblotting, mouse monoclonal anti-HSP90 (610419; BD Biosciences), anti-RFP (M208-3; MBL), and anti-FLAG (F4042; Sigma) antibodies, rabbit polyclonal anti-TEX264 (NBP1-89866; Novus), anti-p62 (PM045; MBL), anti-LC3A (4599s; Cell Signaling), anti-LC3B (4108s; Cell Signaling), anti-GABARAP (13733s; Cell Signaling), and anti-RFP (a gift from Dr. T. Endo, Kyoto Sangyo University) antibodies, and an anti-phosphoserine antibody (ab9332; Abcam) were used as primary antibodies. Anti-mouse (111-035-003; Jackson ImmunoResearch Laboratories, Inc.) and anti-rabbit (111-035-144; Jackson ImmunoResearch Laboratories, Inc.) HRP-conjugated IgGs were used as secondary antibodies. For immunostaining, mouse monoclonal anti-LC3 (M152-3; MBL) was used as a primary antibody. Alexa Fluor 568-conjugated anti-mouse IgG (A-11004; Thermo Fisher Scientific) and Alexa Fluor 660-conjugated anti-rabbit IgG (A-21074; Molecular Probes) antibodies were used as secondary antibodies. A rabbit polyclonal anti-pTEX264 antibody was generated (by ABclonal) against the peptide GASGpSpSFEEL (a.a. 267–276 of TEX264) containing phosphorylated Ser271 and Ser272 and was used for immunoblotting and immunofluorescence microscopy. For transient transfection, Fugene HD (VPE2311; Promega) was used according to the manufacturer’s instructions.

### Preparation of retroviruses

To prepare retroviruses, HEK293T cells were transiently transfected with a retroviral vector together with pCG-VSV-G and pCG-gag-pol (gifts from Dr. T. Yasui, National Institutes of Biomedical Innovation, Health and Nutrition) using Fugene HD. After cells were cultured for 2 days, the supernatant was collected and passed through a 0.45-µm syringe filter unit (SLHV033RB; EMD Millipore).

### Generation of stable cell lines

Cells were cultured with retrovirus and 8 µg/mL polybrene (H9268; Sigma-Aldrich), and stable transformants were selected with blasticidin (022-18713; Wako Pure Chemical Industries) or geneticin (10131; Thermo Fisher Scientific).

### Immunoprecipitation and immunoblotting

Cell lysates were prepared in a lysis buffer (50 mM Tris-HCl, pH 7.5, 150 mM NaCl, 1 mM EDTA, 1% Triton X_-1_00, phosphatase inhibitor cocktail [07574-61; Nacalai Tesque], and protease inhibitor cocktail [EDTA-free]; [03969-34; Nacalai Tesque]). After centrifugation at 17,700 × *g* for 10 min, the supernatants were subjected to immunoprecipitation using anti-FLAG M2 affinity gel (A2220; Sigma-Aldrich). Precipitated immunocomplexes were washed three times in washing buffer (50 mM Tris-HCl, pH 7.5, 150 mM NaCl, 1 mM EDTA, and 1% Triton X_-1_00) and boiled in sample buffer (46.7 mM Tris-HCl, pH 6.8, 5% glycerol, 1.67% sodium dodecyl sulfate, 1.55% dithiothreitol, and 0.02% bromophenol blue). The samples were subsequently separated by sodium dodecyl sulfate-polyacrylamide gel electrophoresis and transferred to Immobilon-P polyvinylidene difluoride membranes (IPVH00010; EMD Millipore). Immunoblotting analysis was performed with the indicated antibodies. Super-Signal West Pico Chemiluminescent Substrate (1856135; Thermo Fisher Scientific) or Immobilon Western Chemiluminescent HRP Substrate (P90715; EMD Millipore) was used to visualize the signals, which were detected on a Fusion System Solo 7S (M&S Instruments, Inc.). Contrast and brightness adjustments and quantification were performed using Fiji software (ImageJ; National Institutes of Health (Schindelin et al., 2012) and Photoshop CS6 (Adobe).

### Immunocytochemistry

Cells grown on coverslips were washed with PBS and fixed with 4% paraformaldehyde in PBS for 10 min at room temperature or 100% methanol for 15 min at −30°C. Fixed cells were permeabilized with 50 µg/mL digitonin (D141; Sigma-Aldrich) in PBS for 5 min, blocked with 3% bovine serum albumin in PBS for 15 min, and incubated with primary antibodies for 1 h. After washing three times with PBS, cells were incubated with Alexa Fluor 568/660-conjugated goat anti-mouse or anti-rabbit IgG secondary antibodies for 1 h. Fluorescence microscopy was performed on a confocal laser microscope (FV3000; Olympus) with a PLANAPO 60×1.4 NA oil-immersion objective lens and captured with FluoView software (Olympus). The number of punctate structures and the ER-autophagosome ratio were determined using Fiji software (ImageJ).

### Cell fractionation

Cells from four 10-cm dishes cultured with 250 nM Torin1 (4247; Tocris Bioscience) and 100 nM bafilomycin A_1_ (B1793; Sigma-Aldrich) for 16 h were harvested and washed twice with ice-cold PBS. The cell pellets were collected after centrifugation at 600 × *g* for 3 min and resuspended in 2 ml ice-cold homogenization buffer (250 mM sucrose, 20 mM Hepes-KOH, pH 7.4, 1 mM EDTA, and complete EDTA-free protease inhibitor). Cells were disrupted by hydrodynamic shearing (27-gauge needle). The homogenized cells were centrifuged twice at 3,000 × *g* for 8 min to remove cell debris and undisrupted cells. The supernatant was diluted with an equal volume of 60% OptiPrep (1114542; CosmoBio) in homogenization buffer. Discontinuous OptiPrep gradients were generated in SW41 tubes (344059; Beckman Coulter) by overlaying the following OptiPrep solutions in homogenization buffer: 2.4 ml of the diluted supernatant in 30% OptiPrep, 1.8 ml in 20%, 2 ml in 15%, 2 ml in 10%, 2.0 ml in 5%, and 2.0 ml in 0%. The gradients were centrifuged at 150,200 × *g* in SW41 Ti rotors (Beckman Coulter) for 3 h, and then 14 fractions (0.8 ml each) were collected from the top. Proteins in each fraction were isolated by precipitation using trichloroacetic acid. The final pellet was suspended in sample buffer and heated at 55°C for 10min.

### In vitro Phosphorylation Assay

Recombinant C-terminally FLAG-Tagged TEX264 was produced by PURExpress in vitro protein synthesis kit (E6800; New England Biolabs). Phosphorylation reaction containing TEX264-FLAG and CK2 (mixture of CK2A1 and CK2B, P6010, NEB) with or without 1 mM CX4549 (S2248; Selleck) were incubated at 30°C for 1h in NEB Buffer (50 mM Tris-HCl, 10 mM MgCl2, 0.1 mM EDTA, 2 mM DTT, and 0.01% Brij 35) and 20 µM ATP. After incubation, the aliquot was suspended with sample buffer and heated at 55°C for 10 min.

### Phosphoproteomic profiling of TEX264

HeLa cells stably expressing TEX264^G280K^-FLAG were cultured with amino acid-free DMEM (048-33575; Wako Pure Chemical Industries) without serum for 1 h and lysed with lysis buffer (50 mM Tris-HCl, pH 7.5, 150 mM NaCl, 1 mM EDTA, 1% NP-40, phosphatase inhibitor cocktail [07574-61; Nacalai Tesque], and protease inhibitor cocktail [EDTA-free]; [03969-34; Nacalai Tesque]). After centrifugation at 17,700 × *g* for 10 min, the supernatants were incubated with anti-FLAG M2 magnetic beads for 3 h at 4°C with gentle rotation. The eluted proteins were enzymatically digested according to a phase-transfer surfactant protocol (Rappsilber *et al*, 2007). Then, 50-μL eluted samples were mixed with 85 μL phase-transfer surfactant buffer. The samples were reduced with 10 mM dithiothreitol at room temperature for 30 min and alkylated with 50 mM of 2-iodoacetamide (804744; Sigma-Aldrich) at room temperature for 30 min. Next, the samples were diluted 5-fold by adding 50 mM NH4HCO3 solution followed by digestion with 1 μg Lysyl Endopeptidase (LysC; 121-05063; Wako Pure Chemical Industries) at 37°C for 4 h. The samples were further digested with 1 μg trypsin at 37°C for 8 h. An equal volume of ethyl acetate acidified with 0.5% TFA was added to the digested samples. After centrifugation at 10,000 × *g* for 10 min twice at room temperature, the aqueous phase containing peptides was collected and dried using an evaporator. The dried peptides were solubilized in 100 μL of 2% acetonitrile and 0.1% TFA. Phosphorylated peptides were enriched using a Titansphere Phos-TiO MP Kit (5010-21281; GL Science) according to the user’s manual. Eluted phosphopeptides were desalted through SDB and GC tips (GL Science). The desalted eluted peptides were then dried using an evaporator.

### Liquid chromatography (LC)-MS/MS analysis

The dried samples were dissolved in 2% acetonitrile and 0.1% TFA and loaded onto the LC-MS system with a Q-Exactive MS instrument (Thermo Fisher Scientific) equipped with a nano HPLC system (Advance UHPLC; Bruker Daltonics) and an HTC-PAL autosampler (CTC Analytics) with a trap column (0.3 × 5 mm, L-column, ODS; Chemicals Evaluation and Research Institute). The samples were separated by a gradient using mobile phases A (0.1% formic acid/H2O) and B (0.1% formic acid and 100% acetonitrile) at a flow rate of 300 nL/min (4% to 32% B for 190 min, 32% to 95% B for 1 min, 95% B for 2 min, 95% to 4% B for 1 min, and 4% B for 6 min) with a home-made capillary column (length of 200 mm and inner diameter of 100 μm) packed with 2-μm C18 resin (L-column2; Chemicals Evaluation and Research Institute). Then, the eluted peptides were electrosprayed (1.8–2.3 kV) and introduced into the MS equipment. Data were obtained using the positive ion mode of data-dependent MS/MS (ddMS^2^) acquisition. Full MS spectra were obtained with a scan range of 350–1800 *m/z* with 70,000 FWHM resolution at 200 *m/z*. MS^2^ spectra were obtained with 17,500 FWHM resolution at 200 *m/z*. For ddMS^2^ acquisition, the 10 highest precursor ions (excluding isotopes of a cluster) above an intensity threshold of 1.7e4 with a charge state from 2+ to 5+ were selected at the 4.0 *m/z* isolation window. A 20-s dynamic exclusion was applied. The obtained raw data were subjected to a database search (UniProt, reviewed mouse database as of September 13, 2018) with Sequest HT algorithm running on Proteome Discoverer 2.2 (Thermo Fisher Scientific). The database was modified to include the TEX264^G280K^ sequence. The parameters for the database search were as follows: peptide cleavage was set to trypsin; missed cleavage sites were allowed to be up to two residues; peptide lengths were set to 6–144 amino acids; and mass tolerances were set to 10 ppm for precursor ions and 0.02 Da for fragment ions. For modification conditions, carbamidomethylation at cysteine was set as a fixed modification. Oxidation at methionine and phosphorylation at serine, threonine, and tyrosine were set as variable modifications. The maximum number of equal modifications per peptide was set to six. A significance threshold of *p* < 0.05 was applied.

### Purification of proteins and peptides

For crystallization and ITC measurement, synthesized TEX264 LIR peptides were purchased from GenScript. Human GABARAP (residues 1–116) and human LC3B (residues 1–120) were prepared as described previously (Mochida *et al*., 2020). Human TEX264 (residues 271–281 with mutation of S272D) fused to the N-terminus of human GABARAP with F3S/V4T mutations was inserted downstream of the human rhinovirus 3C protease recognition site of the pGEX6P-1 vector (Cytiva) and expressed in *Escherichia coli* BL21 (DE3). After the bacteria were cultured at 37°C until OD600 = 1.0, they were supplemented with 100 μM isopropyl-β-D-thiogalactopyranoside and further incubated at 16°C overnight. Cells were collected by centrifugation, resuspended with PBS containing 5 mM EDTA, and sonicated for 10 min. After centrifugation, the recovered supernatants were subjected to GST-accept resins (Nacalai Tesque). The resins were washed three times with PBS and the bound proteins were eluted with 10 mM glutathione and 50 mM Tris, pH 8.0. The eluates were incubated with human rhinovirus 3C protease overnight at 4°C and purified by ion-exchange chromatography using a HiTrap DEAE FF column (Cytiva) with buffers A (20 mM Tris, pH 8.0) and B (20 mM Tris, pH 8.0 and 1 M NaCl) and size-exclusion chromatography using HiLoad 26/60 Superdex 200 prep grade (Cytiva) with 20 mM HEPES, pH 6.8 and 150 mM NaCl. The purified proteins in 20 mM HEPES, pH 6.8 and 150 mM NaCl were concentrated by Centriprep (Millipore) and kept at -80°C until use. Peptides were subjected to size-exclusion chromatography using Superdex peptide 10/300 GL (Cytiva) with 20 mM HEPES, pH 6.8 and 150 mM NaCl and kept at -80°C until use.

### Crystallization and diffraction data collection

All crystallizations were performed through the sitting-drop vapor-diffusion method at 20°C by mixing protein and reservoir solutions in a 1:1 volume ratio. For crystallization of TEX264 LIR^S272D^-GABARAP fusion, the protein was mixed with the reservoir solution consisting of 0.2 M potassium sodium tartrate, 0.1 M sodium citrate, pH 5.6, and 2.0 M ammonium sulfate. For crystallization of GABARAP complexed with TEX264 LIR^p-S271,272^, the protein and peptide were mixed in a 1:1.2 molar ratio and then mixed with the reservoir solution consisting of 30% PEG4000, 0.2 M ammonium acetate, and 0.1 M bis-Tris, pH 5.6. The crystals of TEX264 LIR^S272D^-GABARAP fusion were soaked in the reservoir solution supplemented with 30.7% glycerol and frozen in liquid nitrogen. The flash-cooled crystals were kept in a stream of nitrogen gas at -178°C during data collection. Diffraction data collection was performed using an EIGER X4M detector at beamline BL-1A (KEK) with a wavelength of 1.1000 Å. The diffraction data were indexed, integrated, and scaled using X-ray Detector Software (Kabsch, 2010). The crystals of the GABARAP-TEX264 LIR^p-S271,272^ complex were soaked in the reservoir solution supplemented with 5% glycerol and flash-frozen in a stream of nitrogen gas and kept at -173°C during data collection. Diffraction data collection was performed using an in-house XtaLAB SynergyCustom (Rigaku) that was equipped with a HyPix-6000 detector and MicroMax-007 HF X-ray generator with a wavelength of 1.54178 Å. The diffraction data were indexed and integrated using CrysAlisPro ver. 1.171.40.14a (Rigaku) and merged and scaled using Aimless in the CCP4 program package (Winn *et al*, 2011).

### Structure determination

The structure of the TEX264 LIR^S272D^-GABARAP fusion protein was determined by the molecular replacement method using the Phenix program (Liebschner *et al*, 2019). The crystal structure of GABARAP (PDBID: 1GNU) was used as a search model. Crystallographic refinement was performed using the Phenix and COOT programs (Emsley *et al*, 2010; Liebschner *et al*., 2019). The structure of the GABARAP-TEX264 LIR^p-S271,272^ complex was determined by molecular replacement using MrBUMP (Keegan & Winn, 2008). The crystal structure of GABARAP complexed with AnkB LIR (PDB 5YIR) was used as a search model. Crystallographic refinement was performed using REFMAC5 (Murshudov *et al*, 2011), Phenix, and COOT programs. All structural images in the manuscript were prepared with PyMOL (The PyMOL Molecular Graphics System, Version 2.0 Schrödinger; LLC.).

### ITC

The peptides used in ITC are listed in Fig 4A. ITC experiments were performed using a Microcal iTC200 calorimeter (Malvern Panalytical), with stirring at 1,000 rpm at 25°C. TEX264 peptides were prepared as injection samples from the syringe. The concentration of TEX264 peptides was 0.5 mM. In total, the cell was filled with 200 μL LC3B and GABARAP with a 10-times lower concentration than the syringe samples. The titration involved 18 injections of 2 μL syringe sample at intervals of 120 s into a cell after one injection of 0.4 μL syringe sample. The datasets obtained from the titration of the same syringe sample to the cell filled with buffer were used as reference data for subtraction of the heat of the dilution. MicroCal Origin 7.0 software was used for data analysis. Thermal measurement of the first injection of the syringe samples was removed from the analysis. Thermal titration data were fitted to a single-site binding model, which determines the thermodynamic parameters enthalpy (ΔH), dissociation constant (Kd), and stoichiometry of binding (N). When the fitting was not convergent due to weak interactions, N was fixed to 1.0 or that of the control experiment. The error of each parameter shows the fitting error.

### Doxycycline treatment

HeLa cells stably expressing the ER-phagy probe were cultured with 0.5 μg/mL doxycycline (D3447; Sigma-Aldrich) for 24 h. Thereafter, cells were washed with PBS twice and incubated in DMEM (D6546; Sigma-Aldrich) supplemented with 10% fetal bovine serum (S1820-500; Biowest) and 2 mM L-glutamine (25030-081; Gibco), or amino acid-free DMEM (048-33575; Wako Pure Chemical Industries) without serum followed by confocal microscopy or biochemical analysis.

### Quantification and statistical analysis

Two groups of data were evaluated by an unpaired two-tailed Student’s *t-*test, (ANOVA) followed by Tukey’s multiple comparison test. The data were assumed to have a normal distribution, but this was not formally tested.

### Data availability

Coordinates and structure factors for the GABARAP-TEX264 LIR^p-S271,272^ complex and GABARAP fused to TEX264 LIR^S272D^ have been deposited in the PDB under accession codes 7VEC and 7VED, respectively.

## Acknowledgments

We thank Toshio Kitamura for providing pMXs-IP, Shoji Yamaoka for pMRXIP, Teruhito Yasui for pCG-VSV-G and pCG-gag-pol, Robert A. Weinberg for pCMV-VSV-G, Didier Trono for psPAX2, Toshiya Endo for the anti-RFP antibody, Emi Izumi and Kaori Suzuki for technical assistance with cellular studies and Yuki Ishii for help with protein preparation and crystallization. This study was supported by Exploratory Research for Advanced Technology (ERATO; grant number JPMJER1702 to N.M; JPMJER2001 to H.R.U.) and Core Research for Evolutional Science and Technology (CREST; grant number JPMJCR20E3 to N.N.N.) from the Japan Science and Technology Agency, by JSPS KAKENHI (grant number 19H05707 to N.N.N.; 17K18339 and 20K06637 to A.Y.; JP25221004 to H.R.U.), by HFSP Research Grant Program RGP0019/2018 to H.R.U., and by the Takeda Science Foundation (to N.N.N. and A.Y.).

## Author contributions

H.C. and N.M. designed the project. H.C. performed most cell biology experiments. K.L.O. and H.R.U. performed mass spectrometry. A.Y. and N.N.N. performed ITC analysis and crystal structural analysis. H.C., N.N.N., and N.M. wrote the manuscript. All authors analyzed and discussed the results and commented on the manuscript.

## Conflict of Interest

The authors declare that they have no known competing financial interests or personal relationships that could have appeared to influence the work reported in this paper.

## Expanded View Figure legends

**Figure EV1.**
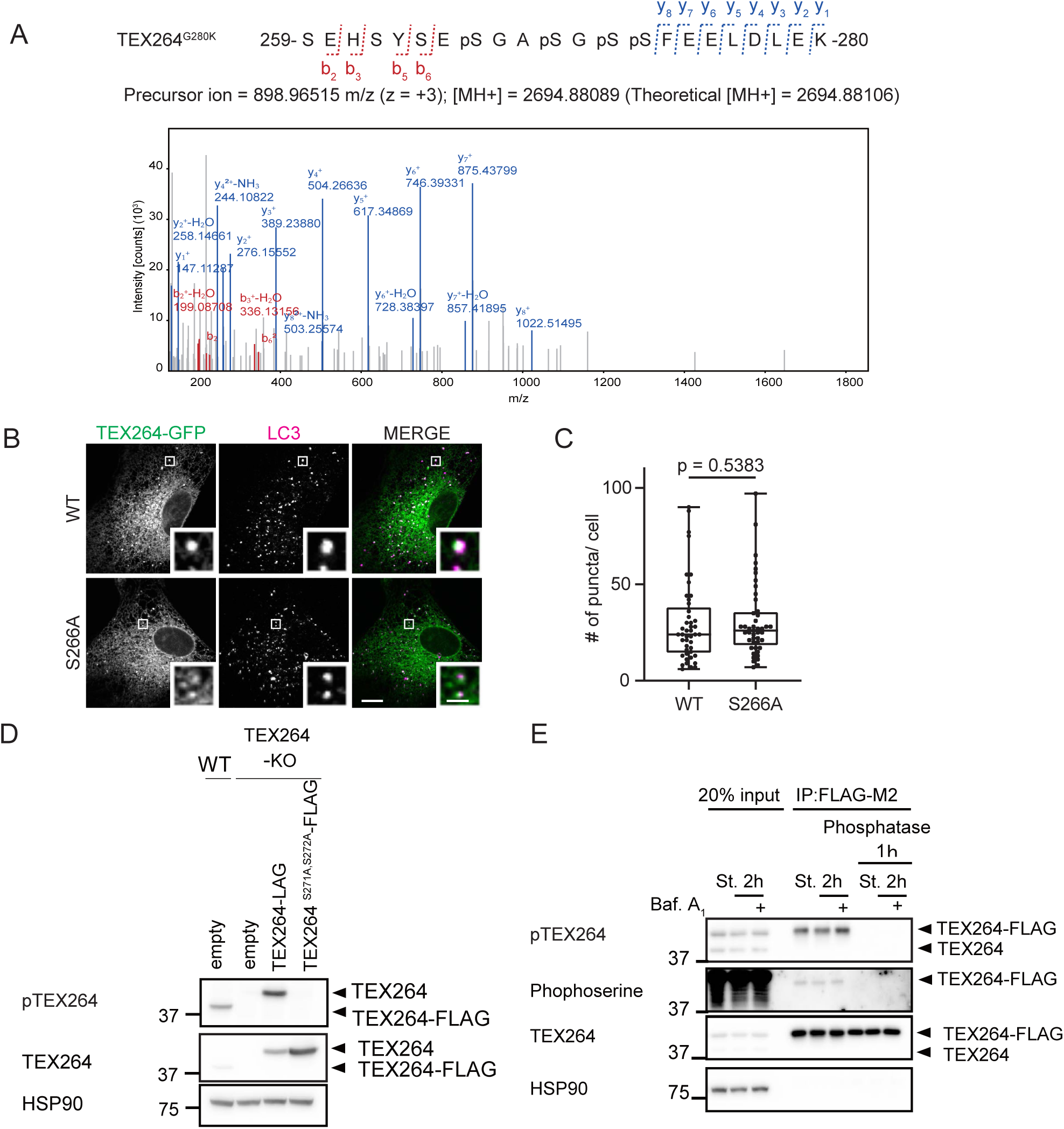
Phosphorylated TEX264 can be detected by an anti-phosphoTEX264 antibody. (A) A representative MS^2^ spectrum of an LIR-containing peptide of TEX264 G280K. B-ions (red), Y-ions (blue) and their neutral losses of water and ammonia are highlighted. (B and C) MEFs stably expressing TEX264-GFP or its S266A mutant were cultured in starvation media and immunostained with an anti-LC3 antibody. Bars: 10 µm and 2 µm (insets) (B). Quantification of the number of TEX264 puncta per cell. Solid bars indicate the medians, boxes the interquartile range (25th to 75th percentile), and whiskers the 0th to 100th percentile. Differences were statistically analyzed by student t-test. Data were collected from 35 cells for each cell type (C). (D) Immunoblotting of the lysates of WT and TEX264-KO HeLa cells stably expressing WT TEX264 or the TEX264^S271A,S272A^ mutant using the indicated antibodies. (E) HeLa cells stably expressing TEX264-FLAG were cultured in starvation media lacking amino acids and serum with or without bafilomycin A_1_ for 2 h. Cell lysates were subjected to immunoprecipitation and treated with or without lambda and alkaline phosphatases for 1 h. Inputs (20% of the lysates) and immunoprecipitates (from 80% of the lysates) were analyzed by immunoblotting using the indicated antibodies.

**Figure EV2.**
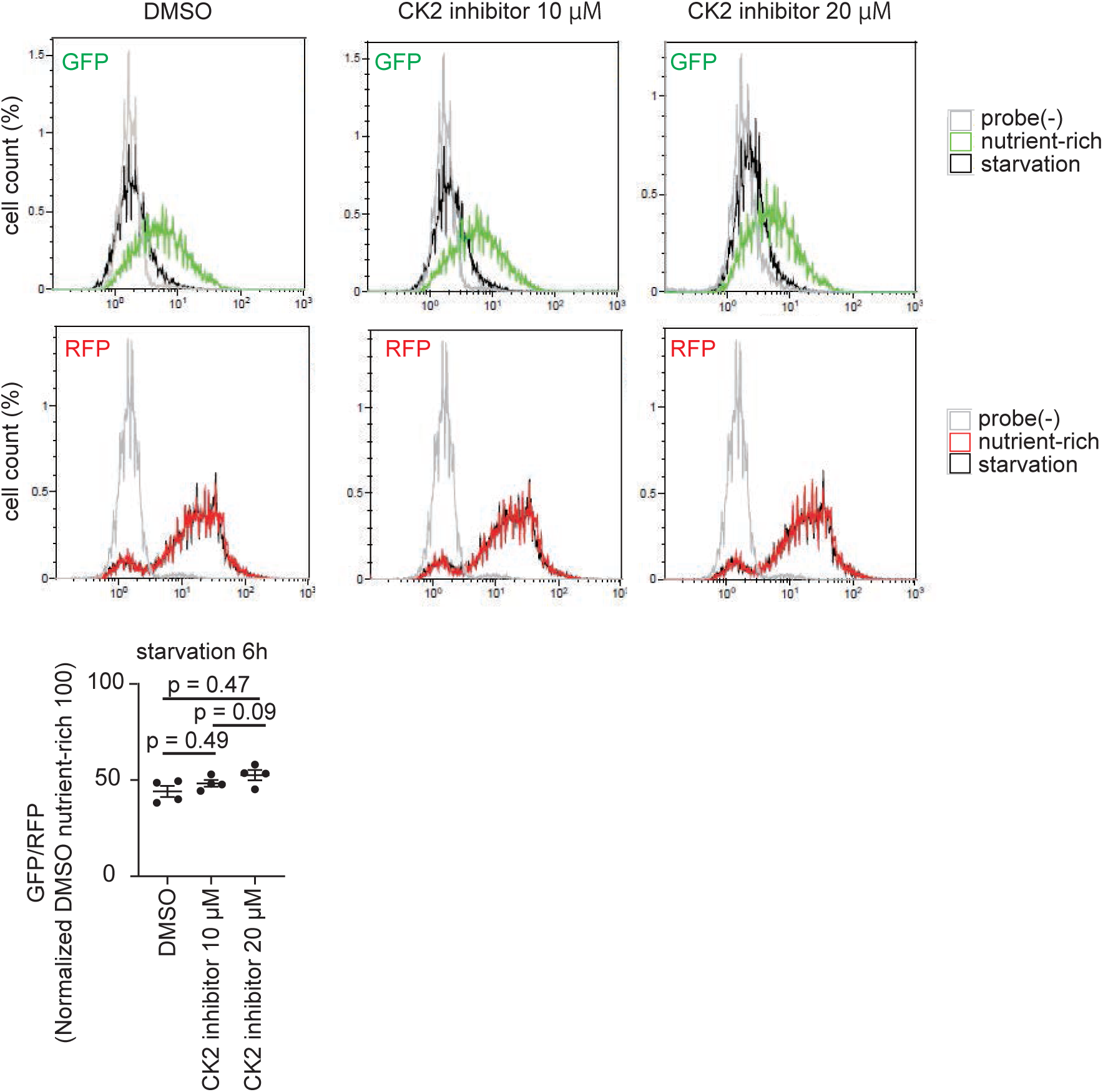
CK2 inhibitor does not affect starvation-induced autophagy. HEK293T cells stably expressing the GFP-LC3-RFP autophagic flux reporter were cultured with 10 µM or 20 µM CK2 inhibitor for 3 h followed by starvation (with the same concentration of inhibitor) for 6 h and analyzed by flow cytometry. The GFP:RFP ratio was quantified. Data represent the mean ± SEM of four independent experiments. Differences were statistically analyzed by one-way ANOVA and Tukey’s multiple comparison test.

**Figure EV3.**
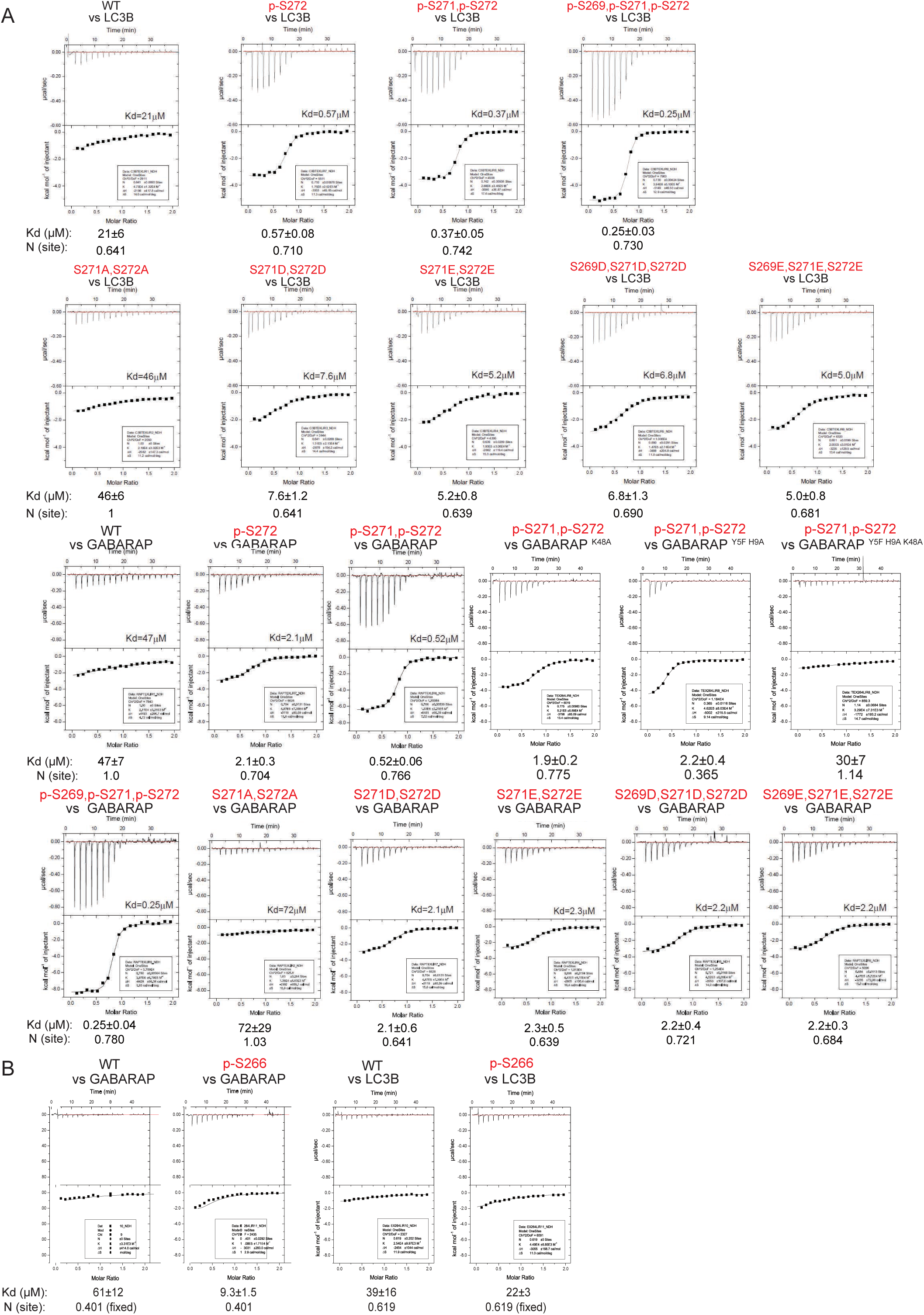
Phosphorylation of TEX264 LIR increases its affinity to ATG8s. Binding affinity of TEX264 LIR peptides (TEX264^269–278^ (A) or TEX264^266–278^ (B)) containing the indicated replacements to LC3B or GABARAP was measured by ITC.

**Figure EV4.**
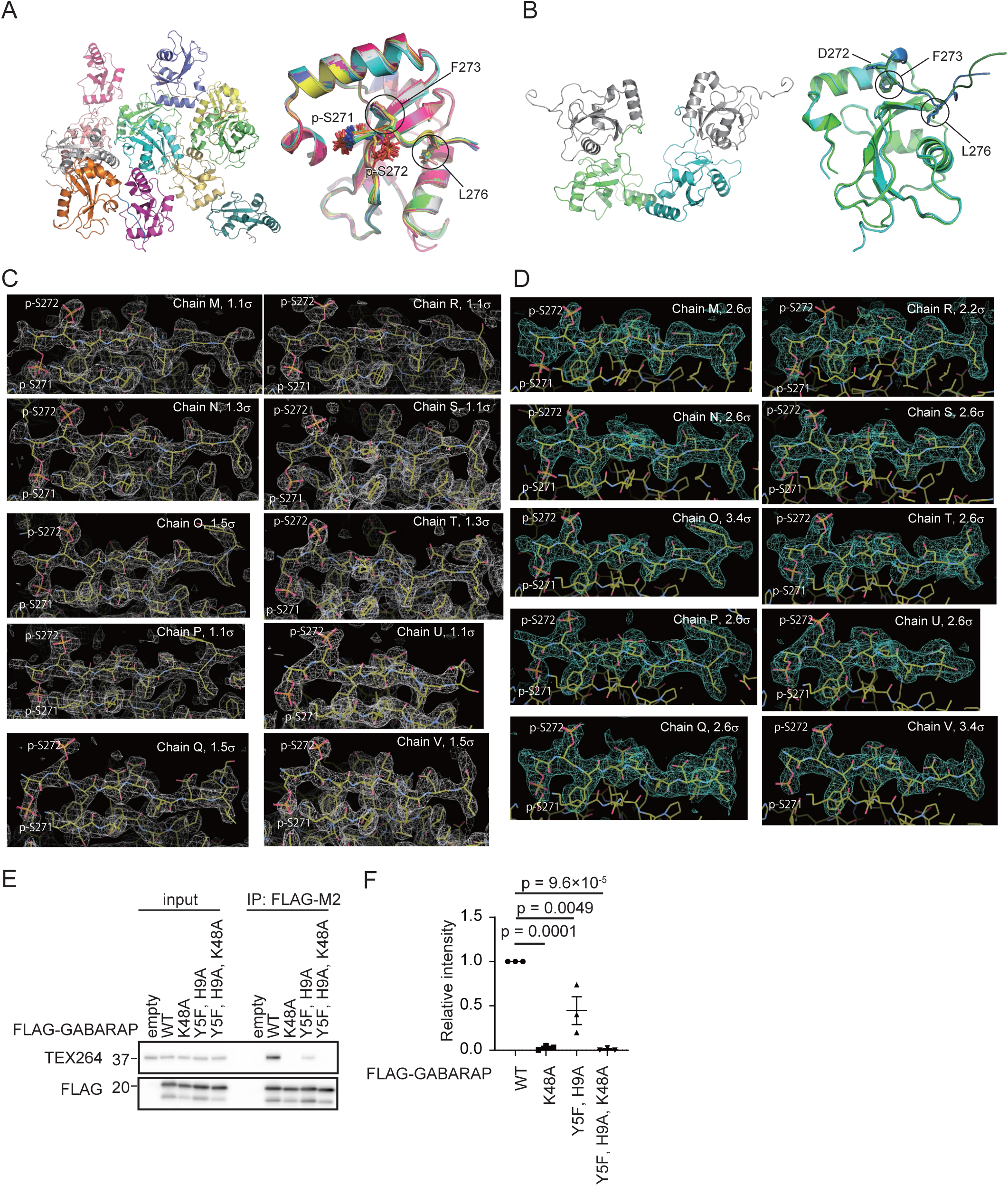
Multiple copies of GABARAP-TEX264 LIR complexes observed in the crystal. (A) Ribbon representation of 12 copies of GABARAP observed in the asymmetric unit of the crystal (left) and their superimposition (right). The side-chains of p-Ser271, p-Ser272, Phe273, and Leu276 are shown with a stick model. (B) Ribbon representation of two copies of GABARAP observed in the asymmetric unit and two symmetrically related GABARAPs in the crystal (left) and their superimposition (right). The side-chains of Asp272, Phe273, and Leu276 are shown with a stick model. (C and D) 2*F*o-*F*c map (C) and simulated annealing *F*o-*F*c omit map (D) of the TEX264 LIR peptides bound to GABARAP together with the structural model, where carbon, nitrogen, oxygen, and sulfur atoms are colored yellow, blue, red, and orange, respectively. Counter levels are indicated at each image. (E and F) HEK293T cells transiently expressing WT or FLAG-GABARAP mutants were subjected to immunoprecipitation (IP). Inputs (20% of the lysates) and immunoprecipitates (from 80% of the lysates) were analyzed by immunoblotting using the indicated antibodies (F). The band intensities in the IP fractions were quantified. Data represent the mean ± SEM of three independent experiments. Differences were statistically analyzed by one-way ANOVA and Tukey’s multiple comparison test (G).

**Figure EV5.**
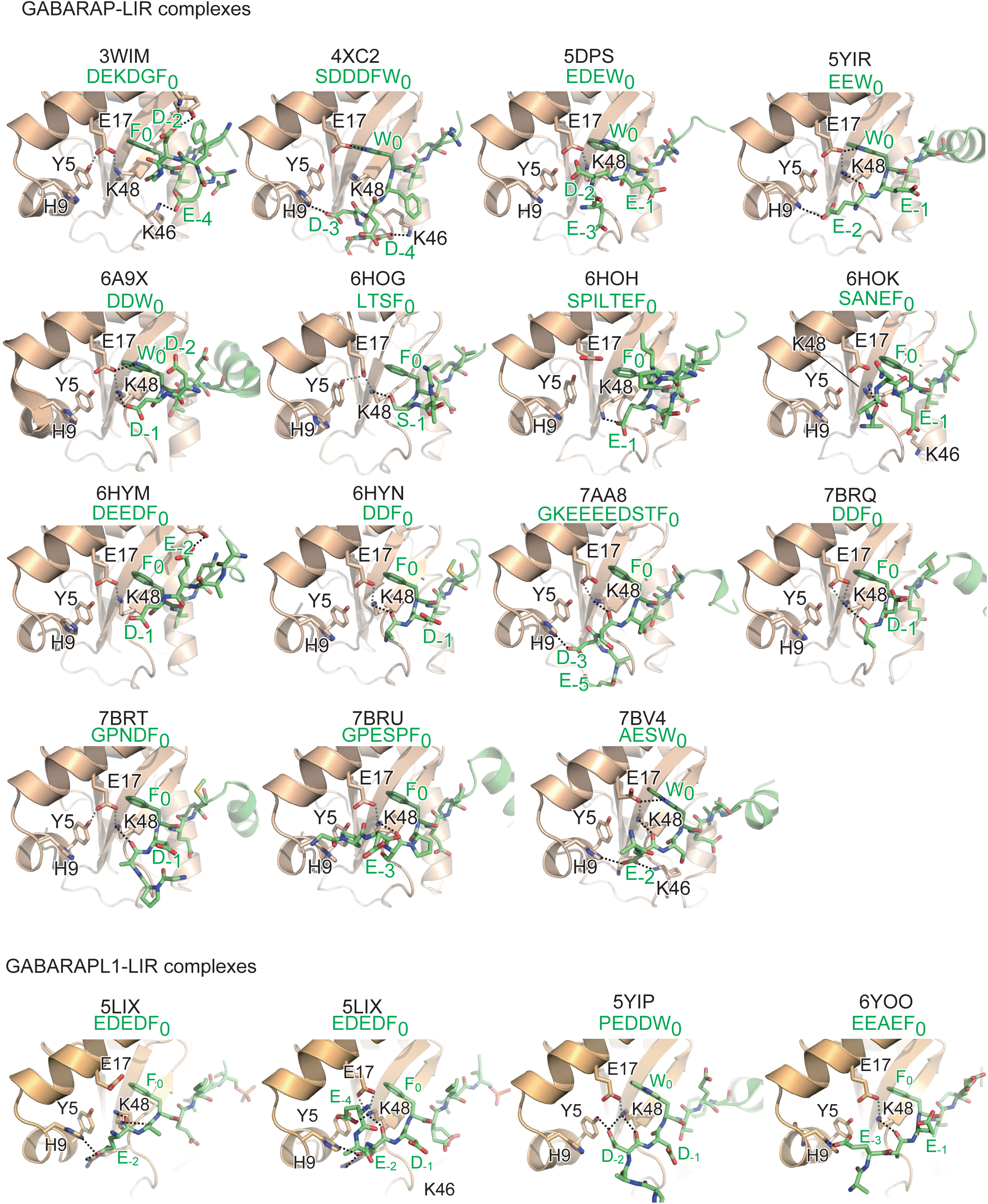
Structural comparison of GABARAP/GABARAPL1-LIR complexes deposited in the PDB. Each structural model is shown in the same orientation as that in Fig 5C. Possible hydrogen bonds are shown as in Fig 5C. Green letters below the PDB code indicate the amino acid sequence in the upstream region of each LIR. Numbers in subscript indicate positions in the LIR sequence.

**Table EV1.** Summary of ITC results.

| Sequence | LC3B |  | GABARAP |  |
| --- | --- | --- | --- | --- |
| | Kd (μM) | N (site) | Kd (μM) | N (site) |
| SGSSFEELDLY | 21±6 | 0.641 | 47±7 | 1.0 (fixed) |
| SGAAFEELDLY | 46±6 | 1.0 (fixed) | 72±29 | 1.03 |
| SGDDFEELDLY | 7.6±1.2 | 0.641 | 2.1±0.6 | 0.728 |
| SGEEFEELDLY | 5.2±0.8 | 0.639 | 2.3±0.5 | 0.696 |
| DGDDFEELDLY | 6.8±1.3 | 0.690 | 2.2±0.4 | 0.721 |
| EGEEFEELDLY | 5.0±0.8 | 0.681 | 2.2±0.3 | 0.684 |
| SGS[pS]FEELDLY | 0.57±0.08 | 0.710 | 2.1±0.3 | 0.704 |
| SG[pS][pS]FEELDLY | 0.37±0.05 | 0.742 | 0.52±0.06 | 0.766 |
| [pS]G[pS][pS]FEELDLY | 0.25±0.03 | 0.730 | 0.25±0.04 | 0.780 |

**Table EV2.**
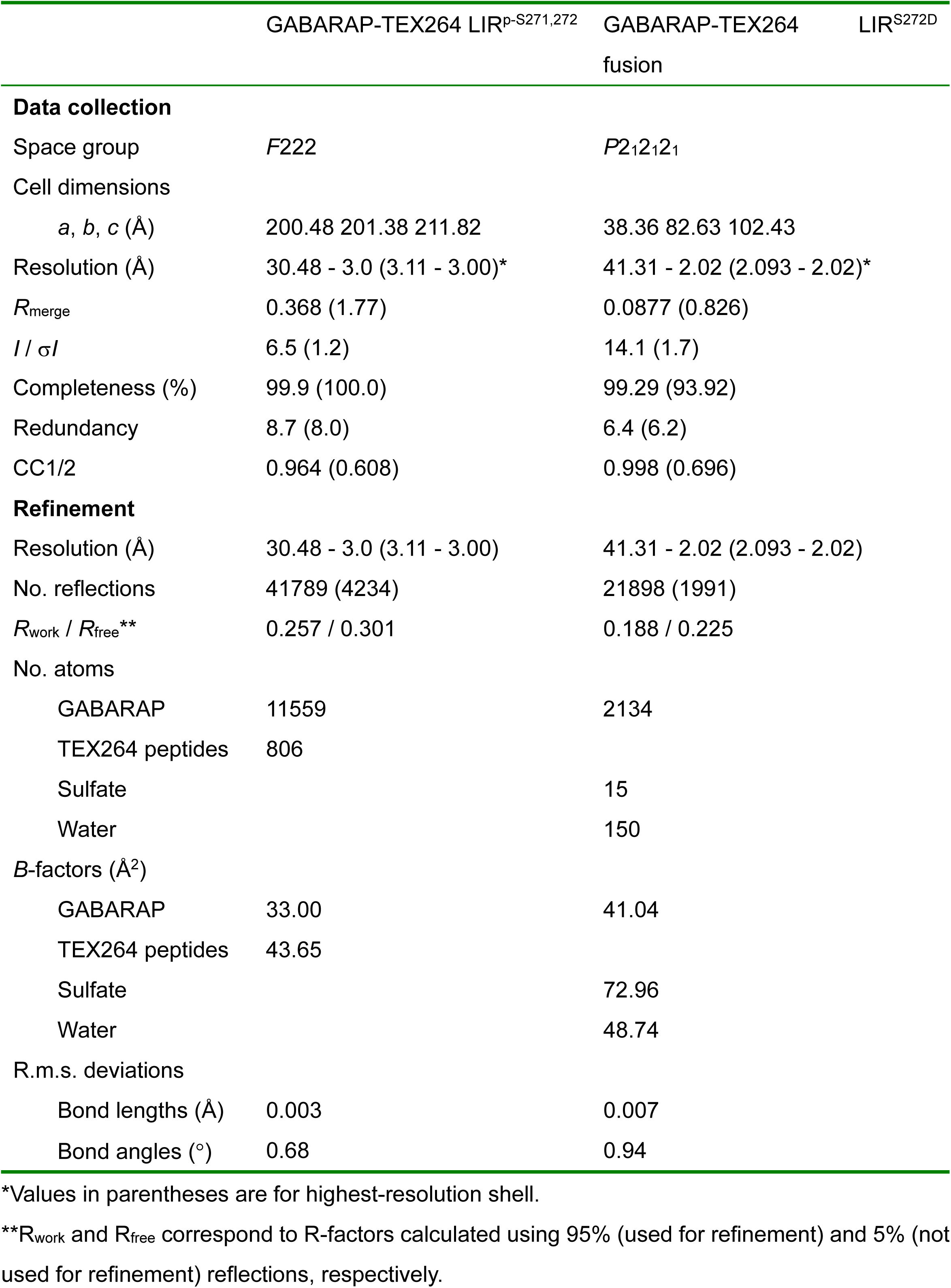
Data collection and refinement statistics.

## Notes

### Competing Interest Statement

The authors have declared no competing interest.

## References

1. An H, Ordureau A, Paulo JA, Shoemaker CJ, Denic V, Harper JW (2019) TEX264 Is an Endoplasmic Reticulum-Resident ATG8-Interacting Protein Critical for ER Remodeling during Nutrient Stress. Mol Cell 74: 891–908.e810

2. Anding AL, Baehrecke EH (2017) Cleaning House: Selective Autophagy of Organelles. Dev Cell 41: 10–22

3. Chino H, Hatta T, Natsume T, Mizushima N (2019) Intrinsically disordered protein TEX264 mediates ER-phagy. Mol Cell 74: 909–921.e906

4. Chino H, Mizushima N (2020) ER-Phagy: Quality Control and Turnover of Endoplasmic Reticulum. Trends Cell Biol 30: 384–398

5. Cooper JA, Sefton BM, Hunter T (1983) Detection and quantification of phosphotyrosine in proteins. Methods Enzymol 99: 387–402

6. Dikic I, Elazar Z (2018) Mechanism and medical implications of mammalian autophagy. Nat Rev Mol Cell Biol 19: 349–364

7. Emsley P, Lohkamp B, Scott WG, Cowtan K (2010) Features and development of Coot. Acta crystallogr D Biol Crystallogr 66: 486–501

8. Fielden J, Wiseman K, Torrecilla I, Li S, Hume S, Chiang SC, Ruggiano A, Narayan Singh A, Freire R, Hassanieh S et al (2020) TEX264 coordinates p97- and SPRTN-mediated resolution of topoisomerase 1-DNA adducts. Nat Commun 11: 1274

9. Gatica D, Lahiri V, Klionsky DJ (2018) Cargo recognition and degradation by selective autophagy. Nat Cell Biol 20: 233–242

10. Gomes LC, Dikic I (2014) Autophagy in antimicrobial immunity. Mol Cell 54: 224–233

11. Hübner CA, Dikic I (2020) ER-phagy and human diseases. Cell Death Differ 27: 833–842

12. Johansen T, Lamark T (2020) Selective Autophagy: ATG8 Family Proteins, LIR Motifs and Cargo Receptors. J Mol Biol 432: 80–103

13. Kabsch W (2010) Integration, scaling, space-group assignment and post-refinement. Acta crystallogr D Biol Crystallogr 66: 133–144

14. Kaizuka T, Morishita H, Hama Y, Tsukamoto S, Matsui T, Toyota Y, Kodama A, Ishihara T, Mizushima T, Mizushima N (2016) An Autophagic Flux Probe that Releases an Internal Control. Mol Cell 64: 835–849

15. Keegan RM, Winn MD (2008) MrBUMP: an automated pipeline for molecular replacement. Acta crystallogr D Biol Crystallogr 64: 119–124

16. Kim J, Kim YC, Fang C, Russell RC, Kim JH, Fan W, Liu R, Zhong Q, Guan KL (2013) Differential regulation of distinct Vps34 complexes by AMPK in nutrient stress and autophagy. Cell 152: 290–303

17. Kirkin V, Rogov VV (2019) A Diversity of Selective Autophagy Receptors Determines the Specificity of the Autophagy Pathway. Mol Cell 76: 268–285

18. Kitamura T, Koshino Y, Shibata F, Oki T, Nakajima H, Nosaka T, Kumagai H (2003) Retrovirus-mediated gene transfer and expression cloning: powerful tools in functional genomics. Exp Hematol 31: 1007-1014

19. Li J, Zhu R, Chen K, Zheng H, Zhao H, Yuan C, Zhang H, Wang C, Zhang M (2018) Potent and specific Atg8-targeting autophagy inhibitory peptides from giant ankyrins. Nat Chem Biol 14: 778–787

20. Liebschner D, Afonine PV, Baker ML, Bunkóczi G, Chen VB, Croll TI, Hintze B, Hung LW, Jain S, McCoy AJ et al (2019) Macromolecular structure determination using X-rays, neutrons and electrons: recent developments in Phenix. Acta crystallogr D Struct Biol 75: 861–877

21. Meggio F, Marin O, Pinna LA (1994) Substrate specificity of protein kinase CK2. Cell Mol Biol Res 40: 401–409

22. Mizushima N (2020) The ATG conjugation systems in autophagy. Curr Opin Cell Biol 63: 1–10

23. Mizushima N, Levine B (2020) Autophagy in Human Diseases. N Engl J Med 383: 1564–1576

24. Mochida K, Yamasaki A, Matoba K, Kirisako H, Noda NN, Nakatogawa H (2020) Super-assembly of ER-phagy receptor Atg40 induces local ER remodeling at contacts with forming autophagosomal membranes. Nat Commun 11: 3306

25. Morita K, Hama Y, Izume T, Tamura N, Ueno T, Yamashita Y, Sakamaki Y, Mimura K, Morishita H, Shihoya W et al (2018) Genome-wide CRISPR screen identifies TMEM41B as a gene required for autophagosome formation. J Cell Biol 217: 3817–3828

26. Murshudov GN, Skubák P, Lebedev AA, Pannu NS, Steiner RA, Nicholls RA, Winn MD, Long F, Vagin AA (2011) REFMAC5 for the refinement of macromolecular crystal structures. Acta crystallogr D Biol Crystallogr 67: 355–367

27. Nakatogawa H (2020) Mechanisms governing autophagosome biogenesis. Nat Rev Mol Cell Biol 21: 439–458

28. Noda NN, Ohsumi Y, Inagaki F (2010) Atg8-family interacting motif crucial for selective autophagy. FEBS letters 584: 1379–1385

29. Palikaras K, Lionaki E, Tavernarakis N (2018) Mechanisms of mitophagy in cellular homeostasis, physiology and pathology. Nat Cell Biol 20: 1013–1022

30. Pinna LA (2002) Protein kinase CK2: a challenge to canons. J Cell Sci 115: 3873–3878

31. Rappsilber J, Mann M, Ishihama Y (2007) Protocol for micro-purification, enrichment, pre-fractionation and storage of peptides for proteomics using StageTips. Nat Protoc 2: 1896–1906

32. Rekha N, Srinivasan N (2003) Structural basis of regulation and substrate specificity of protein kinase CK2 deduced from the modeling of protein-protein interactions. BMC Struc Biol 3: 4

33. Rogov VV, Suzuki H, Fiskin E, Wild P, Kniss A, Rozenknop A, Kato R, Kawasaki M, McEwan DG, Löhr F et al (2013) Structural basis for phosphorylation-triggered autophagic clearance of Salmonella. Biocheml J 454: 459–466

34. Rogov VV, Suzuki H, Marinković M, Lang V, Kato R, Kawasaki M, Buljubašić M, Šprung M, Rogova N, Wakatsuki S, et al (2017) Phosphorylation of the mitochondrial autophagy receptor Nix enhances its interaction with LC3 proteins. Sci Rep 7: 1131

35. Rozenknop A, Rogov VV, Rogova NY, Löhr F, Güntert P, Dikic I, Dötsch V (2011) Characterization of the interaction of GABARAPL-1 with the LIR motif of NBR1. J Mol Biol 410: 477–487

36. Sakurai S, Tomita T, Shimizu T, Ohto U (2017) The crystal structure of mouse LC3B in complex with the FYCO1 LIR reveals the importance of the flanking region of the LIR motif. Acta Crystallogr F Struct Biology Commun 73: 130–137

37. Søreng K, Neufeld TP, Simonsen A (2018) Membrane Trafficking in Autophagy. Int Rev Cell Mol Biol 336: 1–92

38. Sou YS, Waguri S, Iwata J, Ueno T, Fujimura T, Hara T, Sawada N, Yamada A, Mizushima N, Uchiyama Y et al (2008) The Atg8 conjugation system is indispensable for proper development of autophagic isolation membranes in mice. Mol Biol Cell 19: 4762–4775

39. Strickfaden SC, Winters MJ, Ben-Ari G, Lamson RE, Tyers M, Pryciak PM (2007) A mechanism for cell-cycle regulation of MAP kinase signaling in a yeast differentiation pathway. Cell 128: 519–531

40. Suzuki H, Tabata K, Morita E, Kawasaki M, Kato R, Dobson RC, Yoshimori T, Wakatsuki S (2014) Structural basis of the autophagy-related LC3/Atg13 LIR complex: recognition and interaction mechanism. Structure 22: 47–58

41. Thorsness PE, Koshland DE, Jr (1987) Inactivation of isocitrate dehydrogenase by phosphorylation is mediated by the negative charge of the phosphate. J Biol Chem 262: 10422-10425

42. Tsuboyama K, Koyama-Honda I, Sakamaki Y, Koike M, Morishita H, Mizushima N (2016) The ATG conjugation systems are important for degradation of the inner autophagosomal membrane. Science 354: 1036–1041

43. Wild P, Farhan H, McEwan DG, Wagner S, Rogov VV, Brady NR, Richter B, Korac J, Waidmann O, Choudhary C et al (2011) Phosphorylation of the autophagy receptor optineurin restricts Salmonella growth. Science 333: 228–233

44. Winn MD, Ballard CC, Cowtan KD, Dodson EJ, Emsley P, Evans PR, Keegan RM, Krissinel EB, Leslie AG, McCoy A et al (2011) Overview of the CCP4 suite and current developments. Acta crystallogr D Biol crystallogr 67: 235–242

45. Wirth M, Mouilleron S, Zhang W, Sjøttem E, Princely Abudu Y, Jain A, Lauritz Olsvik H, Bruun JA, Razi M, Jefferies HBJ et al (2021) Phosphorylation of the LIR Domain of SCOC Modulates ATG8 Binding Affinity and Specificity. J Mol Biol 433: 166987

46. Zhou J, Tien AC, Alberta JA, Ficarro SB, Griveau A, Sun Y, Deshpande JS, Card JD, Morgan-Smith M, Michowski W et al (2017) A Sequentially Priming Phosphorylation Cascade Activates the Gliomagenic Transcription Factor Olig2. Cell Rep 18: 3167–3177

47. Zhu Y, Massen S, Terenzio M, Lang V, Chen-Lindner S, Eils R, Novak I, Dikic I, Hamacher-Brady A, Brady NR (2013) Modulation of serines 17 and 24 in the LC3-interacting region of Bnip3 determines pro-survival mitophagy versus apoptosis. J Biol Chem 288: 1099–1113

